# Integrative inference of brain cell similarities and differences from single-cell genomics

**DOI:** 10.1101/459891

**Authors:** Joshua Welch, Velina Kozareva, Ashley Ferreira, Charles Vanderburg, Carly Martin, Evan Macosko

## Abstract

Defining cell types requires integrating diverse measurements from multiple experiments and biological contexts. Recent technological developments in single-cell analysis have enabled high-throughput profiling of gene expression, epigenetic regulation, and spatial relationships amongst cells in complex tissues, but computational approaches that deliver a sensitive and specific joint analysis of these datasets are lacking. We developed LIGER, an algorithm that delineates shared and dataset-specific features of cell identity, allowing flexible modeling of highly heterogeneous single-cell datasets. We demonstrated its broad utility by applying it to four diverse and challenging analyses of human and mouse brain cells. First, we defined both cell-type-specific and sexually dimorphic gene expression in the mouse bed nucleus of the stria terminalis, an anatomically complex brain region that plays important roles in sex-specific behaviors. Second, we analyzed gene expression in the substantia nigra of seven postmortem human subjects, comparing cell states in specific donors, and relating cell types to those in the mouse. Third, we jointly leveraged *in situ* gene expression and scRNA-seq data to spatially locate fine subtypes of cells present in the mouse frontal cortex. Finally, we integrated mouse cortical scRNA-seq profiles with single-cell DNA methylation signatures, revealing mechanisms of cell-type-specific gene regulation. Integrative analyses using the LIGER algorithm promise to accelerate single-cell investigations of cell-type definition, gene regulation, and disease states.

## Introduction

The function of the mammalian brain is dependent upon the coordinated activity of highly specialized cell types. Advances in high-throughput single-cell RNAseq (scRNAseq) analysis (Klein et al., 2015; Macosko et al., 2015; Rosenberg et al., 2018; Zheng et al., 2017) have provided an unprecedented opportunity to systematically identify these cellular specializations, across multiple regions (Saunders et al., 2018; Tasic et al., 2016; Zeisel et al., 2018), in the context of perturbations (Hrvatin et al., 2018), and in related species (Hodge et al., 2018; Lake et al., 2016; Tosches et al., 2018). Furthermore, new technologies can now measure DNA methylation (Luo et al., 2017; Mulqueen et al., 2018), chromatin accessibility (Cusanovich et al., 2018), and *in situ* expression (Coskun and Cai, 2016; Moffitt and Zhuang, 2016; Wang et al., 2018), in thousands to millions of cells. Each of these experimental contexts and measurement modalities provides a different glimpse into cellular identity.

Integrative computational tools that can flexibly combine individual single-cell datasets into a unified, shared analysis offer many exciting biological opportunities. For example, systematic comparisons of similar cell types across related brain regions, or homologous tissues from different species, could clarify conserved elements of cell type function, and nominate molecular pathways involved in unique specializations. Additionally, cross-modality analyses—integration of gene expression with spatial information, or epigenomic measurements—could shed important light on the molecular determinants of tissue patterning, and the key mechanisms governing cell-type-specific gene regulation.

The major challenge of integrative analysis lies in reconciling the immense heterogeneity observed across individual datasets. Within one modality of measurement—like scRNA-seq—datasets might differ by many orders of magnitude in the number of cells sampled, or in the depth of sequencing allocated to each cell. Across modalities, datasets may vary widely in dynamic range (gene expression versus chromatin accessibility), direction of relationship (RNA-seq versus DNA methylation), or in the number of genes measured (targeted quantification versus unbiased approaches). To date, the most widely used data alignment approaches (Butler et al., 2018; Haghverdi et al., 2018; Johnson et al., 2007; Risso et al., 2014) implicitly assume that the differences between datasets arise entirely from technical variation and attempt to eliminate them, or map datasets into a completely shared latent space using dimensions of maximum correlation (Butler et al., 2018). However, in many kinds of analysis, both dataset similarities and differences are biologically important, such as when we seek to compare and contrast scRNA-seq data from healthy and disease-affected individuals, or when there are differences cell type composition, with large differences in proportional representation or even cell types missing from some datasets.

To address these challenges, we developed a new computational method called LIGER (**<u>L</u>**inked **<u>I</u>**nference of **<u>G</u>**enomic **<u>E</u>**xperimental **<u>R</u>**elationships). Our approach allows simultaneous, unsupervised discovery of cell types from multiple single-cell experiments, and characterizes both similarities and differences in the properties of these cell types across datasets. We show here that LIGER is extremely robust, enabling the identification of shared cell types across individuals, species, and multiple modalities (gene expression, epigenetic or spatial data), offering a unified analysis of heterogeneous single-cell datasets.

## Results

### Comparing and contrasting single-cell datasets with shared and dataset-specific factors

The intuition behind LIGER is to jointly infer a set of latent factors that represent the same biological “signals” in each dataset, while also retaining the ways in which these signals differ across datasets. These shared and dataset-specific factors can then be used to jointly identify cell types, while also identifying and retaining cell-type-specific differences. LIGER takes as input multiple single-cell datasets, which may be scRNA-seq experiments across different individuals, time points, or species. The inputs to LIGER may even be measurements from different molecular modalities, such as single-cell epigenome data or spatial gene expression data **(Figure 1A).**

**Figure 1:**
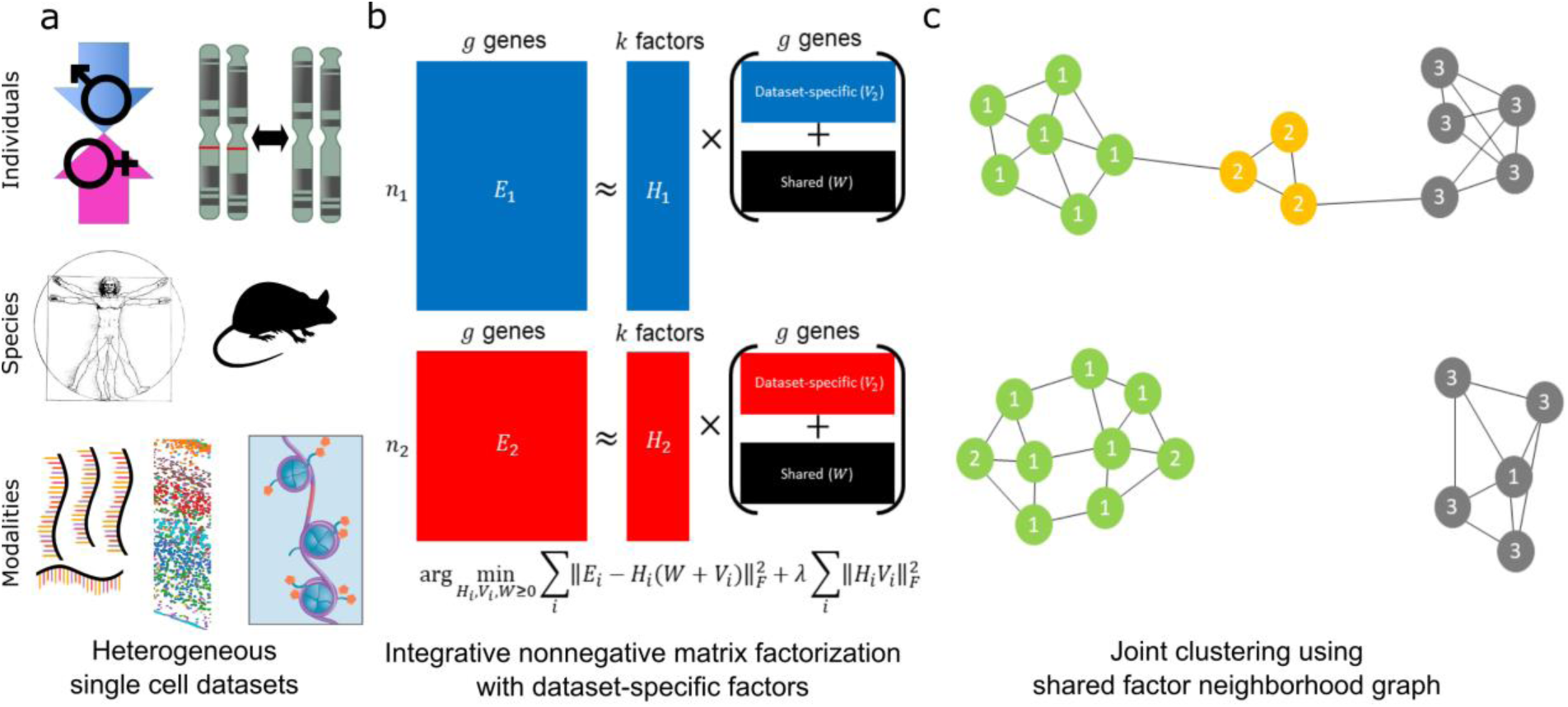
LIGER approach to integration of highly heterogeneous single cell datasets. **(a)** LIGER takes as input two or more datasets, which may come from different individuals, species, or modalities, that share corresponding gene-level features. **(b)** Integrative nonnegative matrix factorization (Yang and Michailidis, 2016) identifies shared metagenes across datasets, as well as dataset-specific metagenes. **(c)** Building a graph in the resulting factor space, based on comparing neighborhoods of maximum factor loadings (**Methods**), allows integration that is highly robust to dataset differences. The schematic shows cells numbered by their maximum factor loadings and connected to their nearest neighbors within each dataset. The shared factor neighborhood graph leverages the factor loading values of neighboring cells to provide additional robustness against noisy factor loadings and prevent the spurious integration of cell types that do not correspond across datasets (such as the yellow cells shown).

LIGER begins by employing integrative non-negative matrix factorization (iNMF) (Yang and Michailidis, 2016) to learn a low-dimensional space in which each cell is defined by one set of dataset-specific factors, and another set of shared factors (**Figure 1B**). Each factor, or metagene, represents a distinct pattern of gene co-regulation, often corresponding to biologically interpretable signals—like the genes that define a particular cell type. The dataset-specific metagenes allow robust representation of highly divergent datasets; for example, in a cross-species scRNA-seq analysis, dataset-specific factors will capture differences in coexpression patterns of homologous genes. The factorization can also accommodate missing cell types by generating a factor with a very large dataset-specific component for each cell type. A tunable term in the factorization objective function penalizes the contributions of dataset-specific metagenes, allowing large or small dataset-specific effects depending on the divergence of the datasets being analyzed. After performing the factorization, we employ a novel clustering strategy that increases the robustness of our results. We first leverage the parts-based nature of the factorization to assign each cell a label based on the factor with the highest loading. We next detect shared clusters across datasets by building a *shared factor neighborhood* graph **(Figure 1C)**, in which we connect cells that have similar neighborhoods of maximum factor loading patterns (**Methods**).

We derived an efficient algorithm that converged in fewer iterations than the standard approach to NMF optimization (**Figures S1A**, **S1B**, and **Methods**), enabling performance that scales well with the size of modern high-throughput single-cell datasets. To aid in selecting the main parameters of the analysis—the number of factors *k* and the tuning parameter λ—we developed heuristics based on factor entropy (measuring the extent to which the information about a cell is encoded in a small number of factors), and dataset alignment (**Methods**). Overall, these heuristics performed well across different analyses (**Figure S1C**), though we have observed that manual tuning can sometimes improve the results. Additionally, we derived novel algorithms for rapidly updating the factorization to incorporate new data or change the k and λ parameters (**Methods** and **Figure S1D**). By using the factors already computed as a starting point for the optimization procedure, we can add new data much more efficiently than simply re-computing the analysis from scratch (**Figure S1E**). So-called “online learning” approaches have proven extremely useful for analyzing massive and continually accumulating data such as internet content, and we anticipate that our approach will be similarly useful for leveraging a rapidly growing corpus of single-cell data.

### LIGER shows robust performance on highly divergent datasets

We assessed the performance of LIGER through the use of two metrics: *alignment* and *agreement*. Alignment (Butler et al., 2018) measures the uniformity of mixing for two or more samples in the aligned latent space by calculating the proportion of nearest neighbors that come from the same versus a different dataset. This metric should be high when datasets share underlying cell-types, and low when attempting to integrate datasets that do not share cognate populations. The second metric, agreement, quantifies the similarity of each cell’s neighborhood when a dataset is analyzed separately, versus when it is analyzed jointly with other datasets by LIGER. A high agreement value indicates that the information about cell types within each dataset is preserved, with minimal distortion, in the joint, integrated analysis.

We calculated alignment and agreement metrics using published datasets (Baron et al., 2016; Gierahn et al., 2017; Saunders et al., 2018), comparing the LIGER analyses to those generated by a recently described method of alignment based on canonical correlation analysis and implemented in the Seurat package (Butler et al., 2018). We first ran our analyses on a pair of scRNA-seq datasets from human blood cells that were prepared by two different technologies and show large-scale, systematic technical variations (Gierahn et al., 2017). Because these datasets measure gene expression from nearly the same cell types, an integrative analysis should yield a high degree of alignment. Indeed, LIGER and Seurat show similarly high alignment statistics (**Figures 2A-C**), with LIGER producing a joint clustering result that is concordant with the published cluster assignments for the individual datasets. LIGER and Seurat performed similarly when integrating human and mouse pancreatic data, with LIGER showing slightly higher alignment **(Figure 2C).**

**Figure 2:**
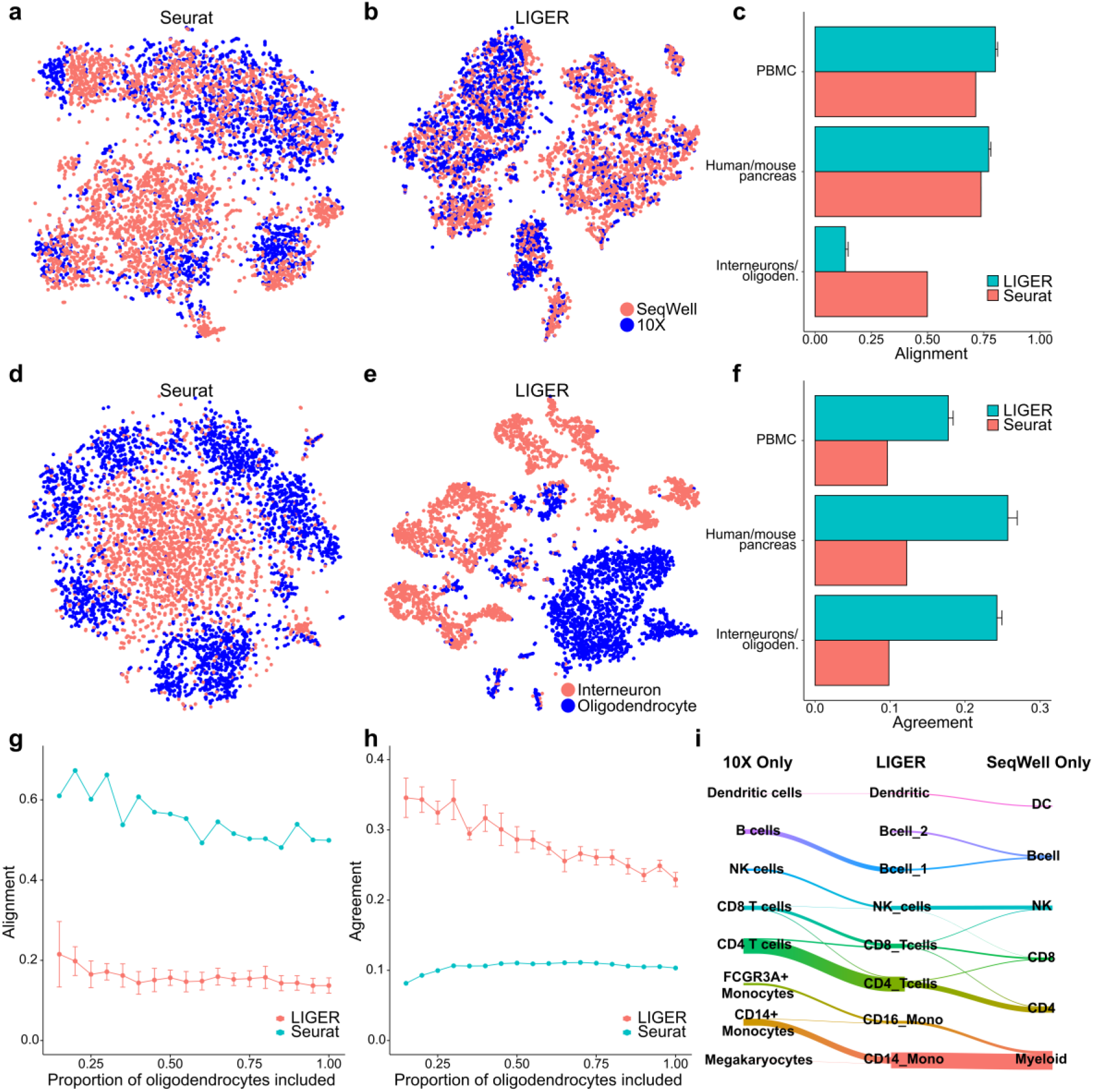
Benchmarking LIGER performance. **(a)** Two-dimensional visualization (t-SNE) of a Seurat/CCA (Butler et al., 2018) analysis of two scRNA-seq datasets prepared from human blood cells. (b) t-SNE visualization of a LIGER analysis of the same dataset analyzed in (a). **(c)** Alignment metrics for the Seurat and LIGER analyses of the human blood cell datasets, human and mouse pancreas datasets, and hippocampal interneuron/oligodendrocyte datasets. Error bars on the LIGER datapoints represent 95% confidence intervals across 20 random initializations. **(d)**-**(e)** t-SNE visualizations of Seurat/CCA (d) and LIGER (e) analyses of 3,212 hippocampal interneurons and 2,524 oligodendrocytes. **(f)** Agreement metrics for Seurat and LIGER analyses of the datasets listed in (c). **(g)**-**(h)** Alignment and agreement for varying proportions of oligodendrocytes mixed with a fixed number of interneurons. **(i)** Riverplot comparing the previously published clustering results for each blood cell dataset with the LIGER joint clustering assignments.

In both analyses, we measured considerably higher agreement in the LIGER analysis (**Figure 2D**), suggesting better preservation of the underlying cell-type architectures when the datasets were brought together in a shared, integrated space. We expected that this advantage should be especially beneficial when analyzing very divergent datasets that share few or no common cell populations. To confirm this advantage of LIGER, we jointly analyzed profiles derived from hippocampal oligodendrocytes and interneurons (Saunders et al., 2018). These cell classes display different developmental trajectories and perform different functions, and thus should not be merged in a joint analysis. The LIGER analysis generated minimal false alignment and demonstrated a good preservation of complex internal substructure (**Figures 2D-F, S2A-C**), allowing clear resolution of many subpopulations, even across considerable changes in dataset proportion **(Figures 2G-H**). To provide an intuitive representation of the sometimes-complex relationships between different clustering results, we used river plots, also called Sankey diagrams, to connect “streams” of cells with the same assignments across cluster analyses. In each of the three analyses described above, the joint LIGER clustering result was highly concordant with the published cluster assignments for the individual datasets (**Figures 2I**, **S2D-E**). Together, these analyses indicate that LIGER shows high sensitivity for discovering common populations without spurious alignment and preserves complex substructure, even when datasets have very few shared signals.

### Interpretable factors unravel complex and dimorphic expression patterns in the bed nucleus

An important application of integrative analysis in neuroscience is to quantify cell-type variation across different brain regions and different members of the same species. To examine LIGER’s performance in these tasks, we analyzed the bed nucleus of the stria terminalis (BNST), a subcortical region comprised of multiple, anatomically heterogeneous subnuclei (Dong and Swanson, 2004) that are implicated in a diverse combination of social, stress-related, and reward behaviors (Bayless and Shah, 2016). To date, single-cell analysis has not yet been deployed on BNST; the robust ability to compare and contrast datasets with LIGER provided an opportunity to clarify how cell types are shared between this structure and datasets generated from related tissues.

We isolated, sequenced, and analyzed 64,000 nuclei enriched for the BNST region (**Figure S3A**, **Methods**). Profiling nuclei rather than whole cells from brain has several advantages, including more faithful representation of cell-type proportions (Habib et al., 2017) and a faster, more systematic isolation approach that avoids transcriptional artifacts associated with cell dissociation (Lacar et al., 2016; Saunders et al., 2018). Initial clustering of these nuclei identified 29,547 neurons; 60.3% of these were localized to BNST by examination of marker expression in the Allen Mouse Brain Atlas (Lein et al., 2007) (**Figure S3B**). Clustering analysis of the BNST-localized neurons revealed 28 transcriptionally distinct populations (**Figure 3A**). In agreement with previous quantification of neurotransmitter identity in this structure (Kudo et al., 2012), 92.4% of the clusters were inhibitory (expressing *Gad1* and *Gad2*), while the remaining 7.6% of clusters were positive for *Slc17a6*, a marker of excitatory cell identity **(Figure S3C**). Examination of markers of these clusters in the Allen Brain Atlas showed that many localized to very distinct BNST substructures, including the principal, oval, rhomboid and anterior commissure nuclei (**Figure S3C** and **S3D**). For example, we identified two very molecularly distinct subpopulations present in the oval nucleus of the anterior BNST (ovBNST) (**Figure 3B**), a structure known to regulate anxiety (Kim et al., 2013), and to manifest a robust circadian rhythm of expression of *Per2* (Amir et al., 2004), similar to the superchiasmatic nucleus (SCN) of the hypothalamus. Interestingly, one of the two ovBNST-localized populations shows specific expression of the VIP receptor *Vipr2*, a gene implicated in circadian rhythm maintenance and male reproduction, suggesting possible functional roles for this specific ovBNST cell type (Dolatshad et al., 2006).

**Figure 3:**
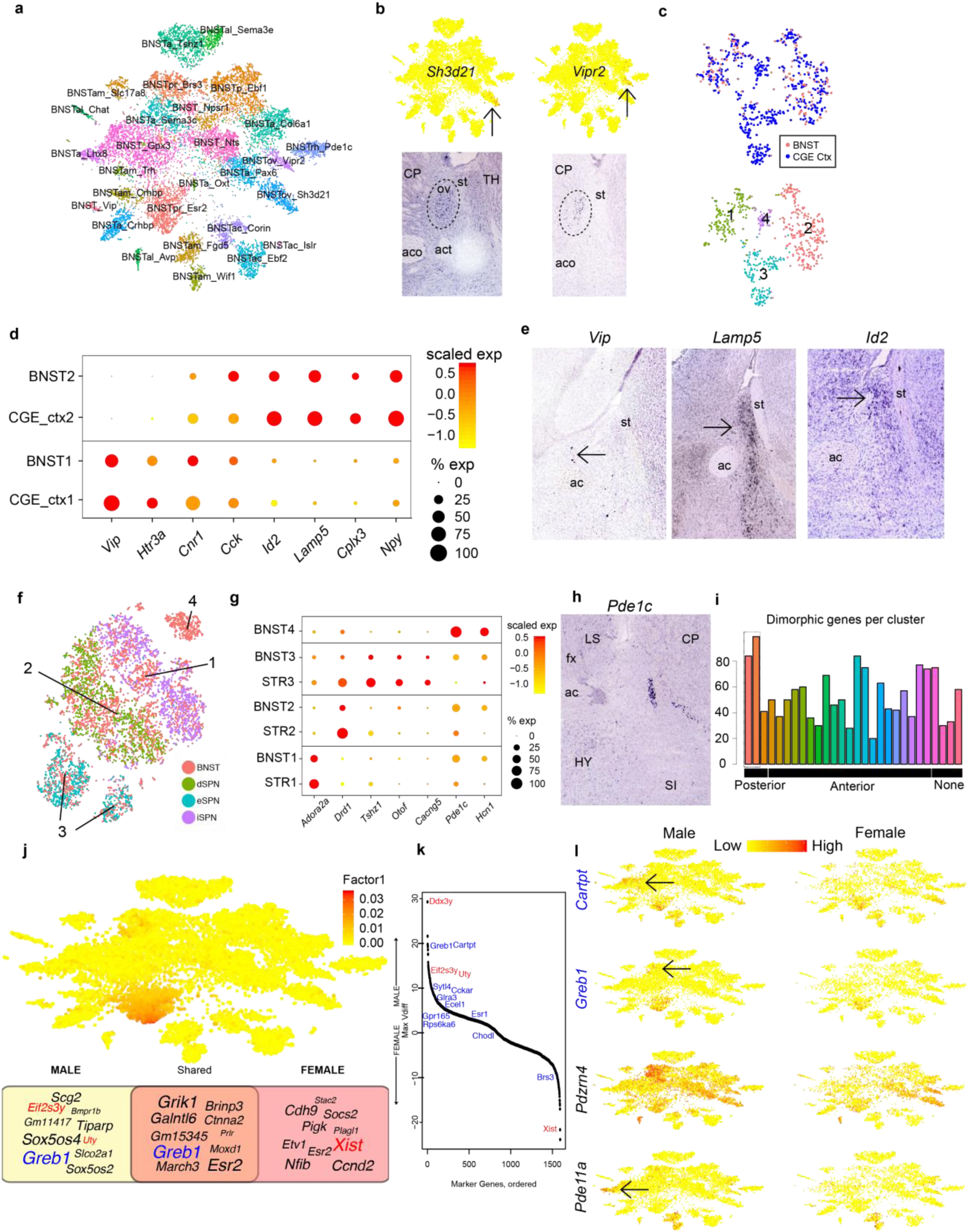
LIGER reveals region-specific and sex-specific cellular specialization in the bed nucleus of the stria terminalis. **(a)** t-SNE visualization of 30,684 bed nucleus neurons analyzed by LIGER, colored by cluster, and labeled by a highly exclusive marker. **(b)** Top, feature plots showing expression of Sh3d21 and Vipr2, in the LIGER BNST analysis. Bottom, sagittal images of ISH for *Sh3d21* and *Vipr2* from the Allen Brain Atlas, showing restricted expression of both markers to the oval nucleus of BNST. **(c)** t-SNE visualization of a LIGER analysis of 139 BNST nuclei in cluster BNST_Vip, and 550 CGE-derived interneurons from frontal cortex (Saunders et al., 2018). On top, points are colored by dataset; bottom shows LIGER-generated cluster assignments. **(d)**. Dot plot showing the relative expression of genes, by dataset, in clusters 1 and 2 of the analysis shown in (c). Each dataset is scaled separately to reconcile differences in sampling between whole cells and nuclei (**Methods**). (**e**) Sagittal ISH images from the Allen Brain Atlas for *Vip*, *Lamp5*, and *Id2*; arrows highlight signal present in the BNST. (**f**) t-SNE visualization of a LIGER analysis of 2,811 BNST nuclei, drawn from three clusters positive for the SPN marker Ppp1r1b (**Figure S3**), and 5,084 striatal SPNs (Saunders et al., 2018). The striatal SPNs are colored according to their previous clustering into three major transcriptional categories (direct, indirect, and eccentric). **(g)** Dot plots showing expression of canonical SPN genes in the clusters defined in (**f**). Markers include those of iSPN identity (*Adora2a*), dSPN identity (Drd1) and the recently described eSPN identity (*Tshz1*, *Otof*, and *Cacng5*), as well as two markers of the BNST-specific cluster 4 (*Pde1c* and *Hcn1*). **(h)** Coronal ISH image of Pde1c from the Allen Brain Atlas, showing strong localization to the rhomboid nucleus of the anterolateral BNST. Abbreviations in ISH images: ac, anterior commissure; fx, fornix; st, stria terminalis; CP, caudate putamen; HY, hypothalamus; LS, lateral septum; SI, substantia innominata. **(i)** Bar plot showing number of dimorphically expressed genes per BNST neuron cluster. Note that the two BNSTpr clusters (leftmost 2 bars) show the most dimorphic genes. **(j)** Cell factor loading values (top) and word cloud plot of top loading dataset-specific and shared genes (bottom) for factor 1, which loads primarily on one of the BNSTpr clusters. The sizes of the gene symbols in the word cloud indicate the strengths of the gene factor loadings. X/Y chromosome genes are colored red, and known dimorphic gene Greb1 is colored blue. Note that Greb1 expression is restricted to the BNSTpr, and thus Greb1 is both a shared cell type marker in males and females and a dimorphically expressed gene. **(k)** Genes ranked by degree of dimorphism; positive values indicate increased expression in males, while negative values indicate increased female expression. Positions of previously validated dimorphic genes and X/Y chromosome genes are indicated in blue and red, respectively. **(l)** Feature plots showing expression patterns of known (*Cartpt* and *Greb1*) and novel (*Pdzrn4* and *Pde11a*) dimorphic genes across BNST neurons.

In addition to identifying cell types that were quite unique to BNST, we found clusters whose marker expression suggested relationships to canonical cell types present in other tissues. One cluster, BNST_Vip, expressed *Vip* and *Htr3a*, two markers of the caudal ganglionic eminence (CGE)-derived interneurons found in cortex and hippocampus. Previous work has shown that at least part of the BNST has embryonic origins in the CGE (Nery et al., 2002), further suggesting that this structure may harbor cell types with similarities to CGE-derived types present in other structures. To examine this possibility, we used LIGER to jointly analyze the 139 nuclei in the BNST_Vip cluster with 550 CGE interneuron cell profiles sampled from a recent large-scale dataset of adult mouse frontal cortex (Saunders et al., 2018). Two clusters in the LIGER analysis showed meaningful alignment between BNST nuclei and cortical CGE cells (**Figure 3C**). Cluster 1 expressed *Vip*, *Htr3a*, *Cck*, and *Cnr1*, likely corresponding to VIP+ basket cells (Rudy et al., 2011) (**Figure 3D**). Examination of *in situ* hybridization images for *Vip* revealed sparse positivity in both the anterior and posterior BNST (**Figure 3E**). The second population, which was *Vip-* negative, expressed *Id2*, *Lamp5*, *Cplx3*, and *Npy*, all markers known to be present in cortical neurogliaform (NG) cells (Tasic et al., 2016) (**Figure 3D**). *In situ* hybridization for two of these markers, *Id2* and *Lamp5*, demonstrated localization of these cells to the principal nucleus (BNSTpr) in the posterior division of BNST (**Figure 3E**). Although, to our knowledge, NG cells have not been described in the BNST before, cells with NG-like anatomy and physiology have been observed within the amygdala (Manko et al., 2012), a structure with related functional roles in cognition and behavior.

SPNs are the principal cell type of the striatum, a structure just lateral to the BNST, but cells expressing the canonical SPN marker, *Pppr1r1b*, have also been documented in multiple BNST nuclei in the anterolateral area (Gustafson and Greengard, 1990). The molecular relationship between striatal SPNs and these BNST cells is not known. We identified three *Ppp1r1b+* populations—two that we annotated as specifically BNST-localized, and a third without BNST-specific localization (**Figures S3B** and **S3D**). To relate these putative SPNs in our dataset to striatal SPNs, we used LIGER to jointly analyze these three clusters (2,811 nuclei, **Figure S3D**), together with 5,084 published SPN profiles from striatum (Saunders et al., 2018). Many of the nuclei from our dataset aligned to clusters 1 and 2 (**Figure 3F**) corresponding to canonical striatal SPNs of the iSPN and dSPN types, respectively. A second population of BNST nuclei aligned to cluster 3, containing the striatal eSPNs, a subset of molecularly defined SPNs that was recently described by scRNA-seq (Saunders et al., 2018). A fourth population, cluster 4, expressed markers localizing it to the rhomboid nucleus of BNST (BNSTrh), suggesting it is a BNST-specific cell type. These results indicate that the BNST likely contains a combination of SPN-like neurons with high homology to striatal SPNs, while also harboring at least one *Ppp1r1b+* population with unique, tissue-specific specializations.

In addition to its high molecular and anatomical diversity, BNST also displays significant sexual dimorphism: in both rodents and primates, it is ∼2-fold larger in males compared to females (Allen and Gorski, 1990; Hines et al., 1992), while several studies have observed gene expression changes in bulk tissue analyses or by *in situ* hybridization for candidate genes. One such study confirmed sexually dimorphic expression in BNST of 12 genes, implicating several of them in regulating sex-specific behaviors (Xu et al., 2012). To better characterize BNST dimorphism at a cell-type-specific level, we jointly analyzed female and male BNST profiles using LIGER, examining the dataset-specific factors to characterize sex-specific differences. We observed that each factor generally loads on only a single cluster; this interpretability allowed us to identify dimorphic expression using the dataset-specific metagenes produced by the factorization. Reassuringly, X- and Y-chromosome genes such as *Xist*, *Eif2s3y* and *Uty* also showed high loading values on dataset-specific factors, reinforcing that these factors were capturing sexually dimorphic gene expression with cell-type-specific resolution. We then used the dataset-specific factor loadings to quantify the number of cell-type-specific dimorphic genes for each cluster (**Methods**).

Our analysis revealed a complex pattern of dimorphic expression involving differences across many individual cell types. Clusters BNSTpr_Brs3 and BNSTpr_Esr2, both localized to the BNST principal nucleus, showed the highest number of dimorphic genes (**Figure 3I**), consistent with previous studies showing the BNSTpr to be particularly dimorphic (Hines et al., 1992; Xu et al., 2012). To illustrate the interpretability of the factorization and the complexity of the dimorphism patterns it reveals, we show the loading pattern and cell-type-specific dimorphic genes derived from a particular factor (**Figure 3J**). A single factor (factor 1) loads strongly on the BNSTpr_Esr2 cluster; among the top dimorphic genes for this factor are *Xist*, *Eif2s3y*, *Uty*, and *Greb1*, one of the 12 previously validated dimorphic genes (Xu et al., 2012). The top loading genes on the shared component of this factor represent cell-type markers, including *Moxd1*, a recently validated marker of BNSTpr (Tsuneoka et al., 2017). *Greb1* itself also loads highly on the shared component of factor 1, reflecting the fact that *Greb1* is a cell-type marker (restricted to the BNSTpr) in both sexes, in addition to showing dimorphic expression. We devised a metric from the LIGER analysis to rank genes by their cell-type-specific dimorphism **(Figure 3K** and **Methods**), flagging genes that are expressed at higher levels in male or female within a specific population. Reassuringly, among the 12 genes previously confirmed to be dimorphic in BNST (Xu et al., 2012), we found that most had high cell-type-specific expression metrics. We also identified new genes with proportional and cell-type-specific dimorphism. The expression patterns of several of the known and new markers that we identified illustrate the complexity of the dimorphism across the many BNST subpopulations (**Figure 3L**).

### Integration of substantia nigra profiles across different human postmortem donors and species

Recent developments in nuclei profiling from archival human brain samples (Habib et al., 2017; Lake et al., 2016) provide an exciting opportunity to comprehensively characterize transcriptional heterogeneity across the human brain, both to understand how specific cell types vary across individual people, and to relate these patterns to large-scale datasets that have been generated from mice. These analyses, however, are challenging. The nuclei are prepared from postmortem samples (fresh *ex vivo* human tissue is only available in very rare circumstances and only for a select few brain regions), which show complex technical variation in gene expression, arising from the influences of many ante- and postmortem variables. The analysis must be capable both of identifying shared cell types across these disparate samples, and of uncovering biologically meaningful changes in cell states, in spite of the significant technical noise.

To explore how well LIGER can deliver an integrated analysis across individual human postmortem samples, we isolated and sequenced 44,274 nuclei derived from the substantia nigra (SN) of seven individuals who were annotated by the brain bank as being neurotypical controls (**Methods**). The SN is a subcortical brain structure with important roles in reward and movement execution, and is known to preferentially degenerate in Parkinson’s disease. Despite considerable inter-individual variation (**Figure S4A**), LIGER successfully delivered an integrated analysis in which each of the cell-type substituents of the SN was accurately aligned across datasets **(Figure 4A).** Specifically, we identified 24 clusters that included all known resident cell classes: astrocytes, fibroblasts, mural cells, microglia, neurons (including TH+ dopaminergic neurons and multiple inhibitory types), oligodendrocytes, and oligodendrocyte progenitor cells (polydendrocytes) **(Figure 4B-C).**

**Figure 4:**
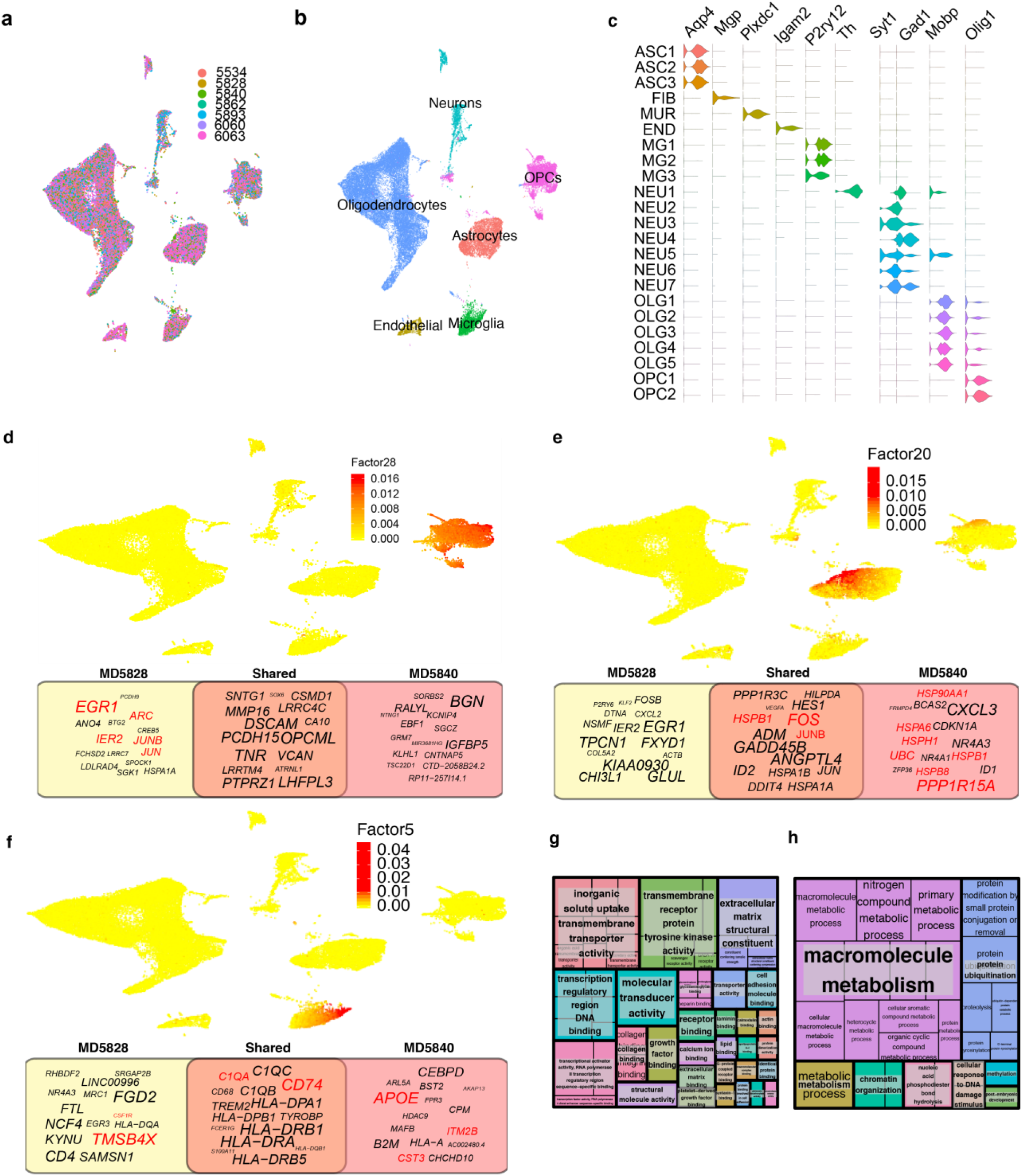
LIGER allows analysis of substantia nigra across individuals and species. **(a)-(b)** Two-dimensional UMAP representation of a LIGER analysis of 44,274 nuclei derived from the substantia nigras of 7 human donors, colored by donor **(a)** and major cell class **(b). (c)** Violin plots showing expression of marker genes across the 25 human SN populations identified by two rounds of LIGER analysis. **(d)-(f)** UMAP plots showing cell factor loading values (top) and word cloud plots (bottom) for a factor corresponding to an acutely activated polydendrocyte state **(d)**, an activated microglia state **(e)** and a reactive astrocyte state **(f).** In the word clouds, the size of gene text in the word clouds represents the relative contribution of that gene to the shared or dataset-specific metagenes. Genes in the yellow box load on the metagene specific to tissue donor MD5828, who suffered a traumatic brain injury at the time of death; genes in the pink box load highly on the metagene specific to tissue donor MD5840, who was diagnosed with cerebral amyloid angiopathy. Genes in the orange box load on the shared metagene common to all datasets; genes mentioned in the text are highlighted red. **(g)** GO terms enriched in homologous genes with strong expression correlation across substantia nigra clusters in the LIGER comparative analysis of human and mouse. **(h)** GO terms enriched in homologous genes with weak expression correlation.

Our integrated analysis offered an opportunity to assess the variation of cell-type-specific expression across individual tissue donors. Glial activation is an important hallmark and driver of many brain diseases, including neurodegeneration and traumatic brain injury. To uncover datasets with atypical glial expression patterns, we searched the dataset-specific metagenes of glial cell types for evidence of altered states. Examination of the polydendrocyte-specific factor 28 showed that subject MD5828 had high expression of immediate early genes (**Figure 4D**), including *FOS*, *ARC, IER2,* and *EGR1,* consistent with an acute injury (Dimou et al., 2008). Although this subject was coded as a control, examination of cause of death revealed a very strong likelihood of brain trauma (**Methods**). In addition, the MD5828-specific metagene for factor 5, a microglia-specific factor, showed high loadings of *TMSB4X* and *CSF1R*, both of which play important roles in the acute response to traumatic brain injury (Luo et al., 2013; Xiong et al., 2012). By contrast, in subject 5840, the dataset-specific loadings on the microglial factor 5 included genes known to be upregulated in microglia in response to amyloid deposition, such as *APOE* and *TREM2* (**Figure 4E**). Review of this subject’s postmortem report revealed a histological diagnosis of moderate-severity cerebral amyloid angiopathy (CAA), a disease in which amyloid deposits within the walls of CNS vasculature. Intriguingly, two of the three genes known to cause hereditary CAA (Biffi and Greenberg, 2011), *CST3* and *ITM2B*, were also strong contributors to the MD5840-specific factor 5 metagene. In an astrocyte-specific factor (factor 20), subject MD5840 showed remarkable upregulation of multiple genes involved in protein misfolding response (**Figure 4F**), including *UBC*, *HSPA6*, *HSPB1*, *HSPH1*, *HSP90AA1*, *HSPB8*, *HSPD1* and *PPP1R15A*, a master regulator of the protein misfolding response pathway (Tsaytler et al., 2011). Several of these genes have been directly implicated in the reactive glial response to amyloid accumulation in cerebral vessels (Bruinsma et al., 2011).

A deeper understanding of cell types often arises from comparisons across species. For example, cell types with similar morphologies and even functions have occasionally been found, through comparative molecular analyses, to have surprisingly divergent evolutionary origins (Arendt et al., 2016). We therefore used LIGER to compare the SN across species, integrating our newly generated human data with a recently published single-cell dataset from the mouse SN (Saunders et al., 2018). This is an especially challenging analysis: first, mice and humans are separated by 96 million years of evolution, causing widespread deviations in gene regulation within homologous cell types. Second, the human data derive from frozen, post-mortem nuclei, while the mouse data were prepared from whole dissociated cells extracted from fresh, perfused brain tissue. Despite these challenges, LIGER was able to successfully identify both the corresponding broad cell classes across species, and subtler cell types within each class after a second round of analysis **(Figures S4B-S4F**). In our subanalysis of the neurons, we found that LIGER avoided false positive alignments of human profiles to mouse cell types outside the dissection zone of the human tissue (**Figures S4G** and **S4H**), reinforcing LIGER’s ability to perform very noisy, complex integrative analyses across species. Overall, we observed strong concordance between mouse and human cell clusters, consistent with another recent comparative single-cell analysis of mouse and human cortex (Hodge et al., 2018). The only two clear species-specific populations in our analysis could be attributed to differences in dissection between human and mouse (**Figures S4F-H**).

Understanding how expression of homologous genes within the SN differs across species could give insight into differences in how these genes function within the tissue. Using the set of joint clusters assigned by LIGER, we aggregated gene expression across each cluster, for each species, and computed pairwise Pearson correlation values for each mouse gene and its human homolog. We performed a gene ontology (GO) term enrichment analysis to evaluate whether genes with the highest and lowest correlation share any functional relationships. We found that the homologous gene pairs with high expression correlation were enriched for GO terms related to brain cell identity and basic molecular functions, including ion channels, transcription factors, transmembrane receptors, and extracellular matrix structural components (**Figure 4G**). In contrast, the least correlated homologous gene pairs were enriched for basic metabolic processes, including macromolecule metabolism, protease activity, and DNA repair (**Figure 4H**). Intriguingly, genes involved in chromatin remodeling (sharing the GO function “chromatin organization”) also showed less expression conservation, hinting at species differences in epigenetic regulation.

### Integrating scRNA-seq and *in situ* transcriptomic data locates frontal cortex cell types in space

Spatial context within a complex tissue is an important aspect of a cell’s identity. Knowing where a cell is located can give important insights into functions within a tissue and signaling interactions with nearby cells. Additionally, spatially resolved transcriptomic data provide an invaluable opportunity to confirm the significance of cell clusters defined from scRNA-seq data, because transcriptomic distinctions that also correspond to unique spatial localization patterns likely represent biologically relevant differences. Integrated analysis of spatial transcriptomic and scRNA-seq data using LIGER could offer two potential advantages compared to separate analyses of the two data modalities: (1) assigning spatial locations to cell clusters observed in data from dissociated cells; and (2) increasing the resolution for detecting cell clusters from the *in situ* data.

To explore how well LIGER can successfully integrate spatial and whole-transcriptomic datasets, we jointly analyzed frontal cortex scRNA-seq data prepared by Drop-seq (Saunders et al., 2018) and *in situ* spatial transcriptomic data from the same tissue generated by STARmap (Wang et al., 2018). These two datasets differ widely in many respects, including in number of cells (71,000 scRNA-seq vs. 2500 STARmap) and genes measured per cell (scRNA-seq is unbiased, while STARmap assays only selected markers). Nevertheless, LIGER was able to correctly define joint cell populations across the datasets (**Figure 5A-B**), with expression of key marker genes confirming the correspondence of cells across these different modalities **(Figure 5C).** Reassuringly, only one population in the scRNA-seq data was dataset-specific, corresponding to cells from the claustrum, an anatomical structure that was not included in the tissue assayed by STARmap. Because the spatial locations of the STARmap profiles are known, our integrated analysis using LIGER allowed us to spatially locate each of the jointly defined populations (**Figure 5D**). Our results accorded well with the known spatial features of the mouse cortex, including meninges and sparse layer 1 interneurons at the surface, excitatory neurons organized in layers 2-6, and oligodendrocyte-rich white matter below the cortex (**Figure 5D**). One replicate of the STARmap data also showed a chain of co-localized endothelial cells running through the cortex, presumably a contiguous segment of vasculature **(Figure 5D).** The success of this integrative analysis is especially noteworthy given the very different global distributions of gene expression values in the scRNA-seq data compared to the STARmap data—the scRNA-seq data are sparse, with many zero values, while the STARmap data show no zero-inflation (**Figure 5E**).

**Figure 5:**
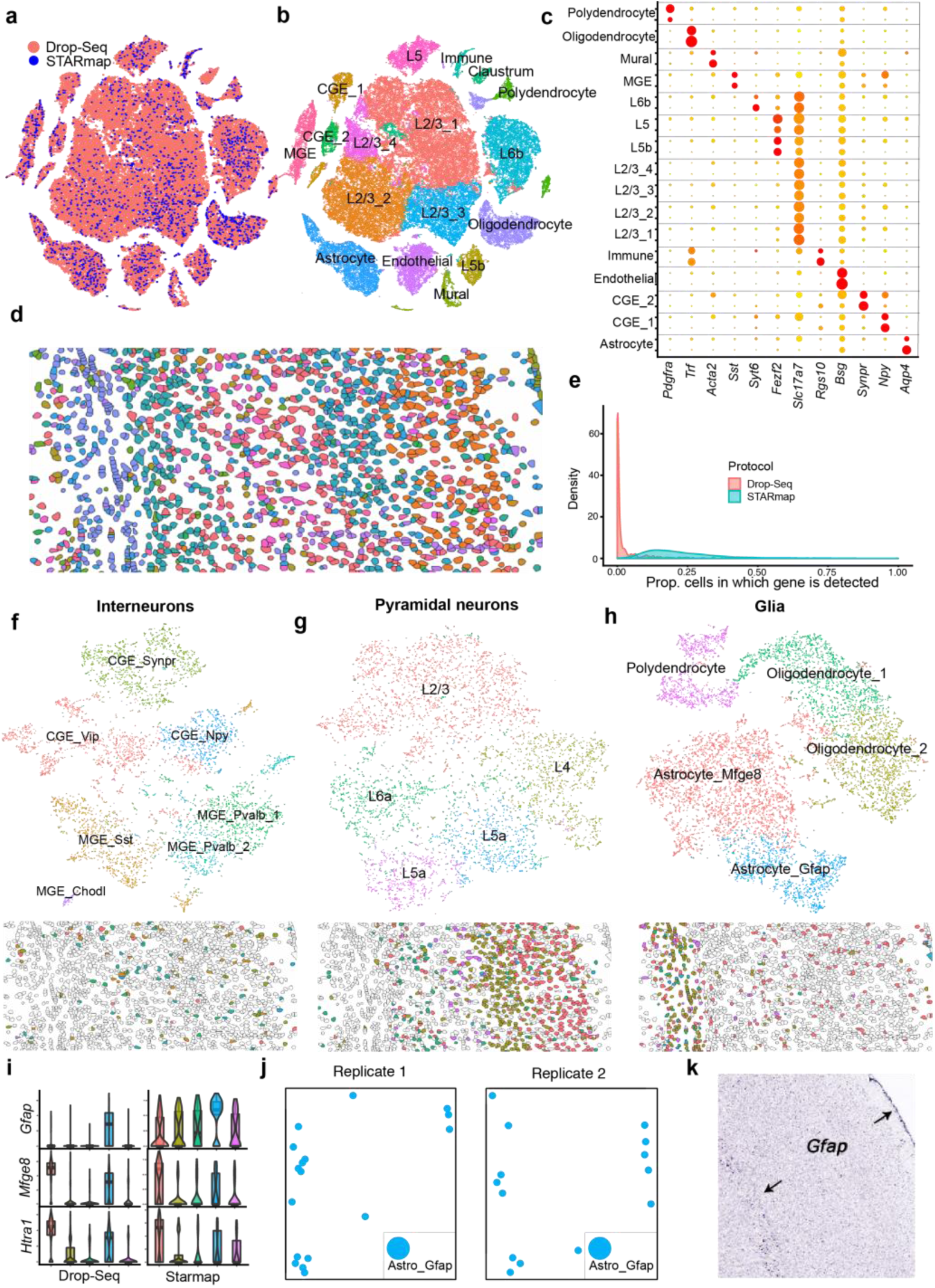
Locating cortical cell types in space using scRNAseq and STARmap. **(a)-(b)** t-SNE visualizations of a LIGER analysis of 71,000 frontal cortex scRNAseq profiles prepared by Drop-seq (Saunders et al., 2018) and 2,500 cells profiled by STARmap (Wang et al., 2018) colored by technology **(a)** and LIGER cluster assignment **(b)**. Labels in (b) derive from the published annotations of the Drop-seq dataset. **(c)** Dot plot showing marker expression for STARmap cells (top line of each gene) and Drop-seq cells (bottom line) across LIGER joint clusters. **(d)** Spatial locations of STARmap cells colored by LIGER cluster assignments. **(e)** Density plot showing proportion of cells in which each gene is detected for the Drop-seq (red) and STARmap (blue) datasets. **(f)-(h)** t-SNE plots and spatial locations for LIGER subclustering analyses of interneurons **(f)**, pyramidal neurons **(g)** and glia **(h)**. **(i)** Violin plots of marker genes for two astrocyte populations identified in subclustering analysis of glia. **(j)** Spatial coordinates for *Gfap* expressing astrocyte populations (two STARmap replicates shown). **(k)** *Gfap* staining data from the Allen Brain Atlas showing localization of Gfap to both meninges and white matter layer below cortex.

Incorporating the scRNA-seq data, which contained whole-transcriptomic measurements from more than 28 times as many cells as the STARmap data, allowed us to identify cell populations with greater resolution. We performed a second round of clustering on subsets of cells that were grouped together in the coarser first-round clustering, of inhibitory interneurons (**Figure 5F**), excitatory neurons (**Figure 5G**), and glia (**Figure 5H**). We identified 7 interneuron clusters and 5 glial clusters compared to 4 and 2 clusters, respectively, in the initial STARmap analysis. These additional populations accorded well with cell-type distinctions defined in the original scRNA-seq analysis. The 5 glial clusters we identified included two astrocyte clusters, polydendrocytes, and two clusters of oligodendrocytes (Wang et al., 2018). The two astrocytic subpopulations expressed patterns of marker genes consistently between both the scRNA-seq and STARmap datasets (**Figure 5F**). The larger population expressed high levels of *Mfge8* and *Htra1*, while the second population showed high expression of *Gfap* (**Figure 5F**). We found that the *Gfap*-expressing astrocyte population is located outside the cortical gray matter, in both the meningeal lining and the white matter below layer 6 (**Figure 5G, 5H**), consistent with a more fibrous identity. In contrast, the larger second population of astrocytes was spread uniformly throughout the cortical layers, consistent with a protoplasmic phenotype. Identifying the localization pattern of the *Gfap*-expressing astrocyte population also helped to clarify a finding from our human-mouse substantia nigra analysis (**Figure S4E)**, suggesting that this same *Gfap*-expressing population is likely missing from the human data because of dissection differences. These results show the clarifying power of jointly leveraging large-scale scRNA-seq and *in situ* gene expression data for defining cell types in the brain.

### LIGER defines cell types using both single-cell transcriptome and single-cell DNA methylation profiles

Methods of measuring epigenomic regulation in individual cells provide a new means of characterizing cellular heterogeneity beyond gene expression, as well as several exciting but relatively unexplored analytical opportunities if they can be linked with scRNA-seq data. First, the congruence between epigenome-defined cell populations and scRNA-seq-defined populations remains unclear; it is unknown whether clusters defined from gene expression reflect epigenetic distinctions and vice versa. Second, integrating single-cell epigenetic and transcriptomic data provides an opportunity to study the mechanisms by which epigenetic information regulates gene expression to determine cell identity. Finally, integrating single-cell epigenetic profiles with scRNA-seq data may improve sensitivity and interpretability compared to analyzing the epigenetic data in isolation, since scRNA-seq technology can offer greater throughput and capture more information per cell.

To investigate these possibilities, we performed an integrated analysis of two single-cell datasets prepared from mouse frontal cortical neurons: one that measured gene expression using Drop-seq (55,803 cells) (Saunders et al., 2018) and another that measured genome-wide DNA methylation (3,378 cells) (Luo et al., 2017). We reasoned that, because gene body methylation is generally anticorrelated with gene expression (Mo et al., 2015), reversing the direction of the methylation signal would allow joint analysis. Although multiple epigenetic mechanisms regulate gene expression, and the relationship between expression and methylation for individual genes may be weak or even reversed, a generally inverse relationship between methylation and expression across many genes should enable joint identification of cell types across modalities. The number of cells, large imbalance in dataset sizes, different modalities, and inverse relationship between methylation and expression pose significant challenges to data integration. Despite these challenges, LIGER successfully integrated the datasets, producing a joint clustering that identified the dominant neuronal cell types of the frontal cortex and accorded very well with the published analyses of each dataset (**Figure 6A-C**).

**Figure 6:**
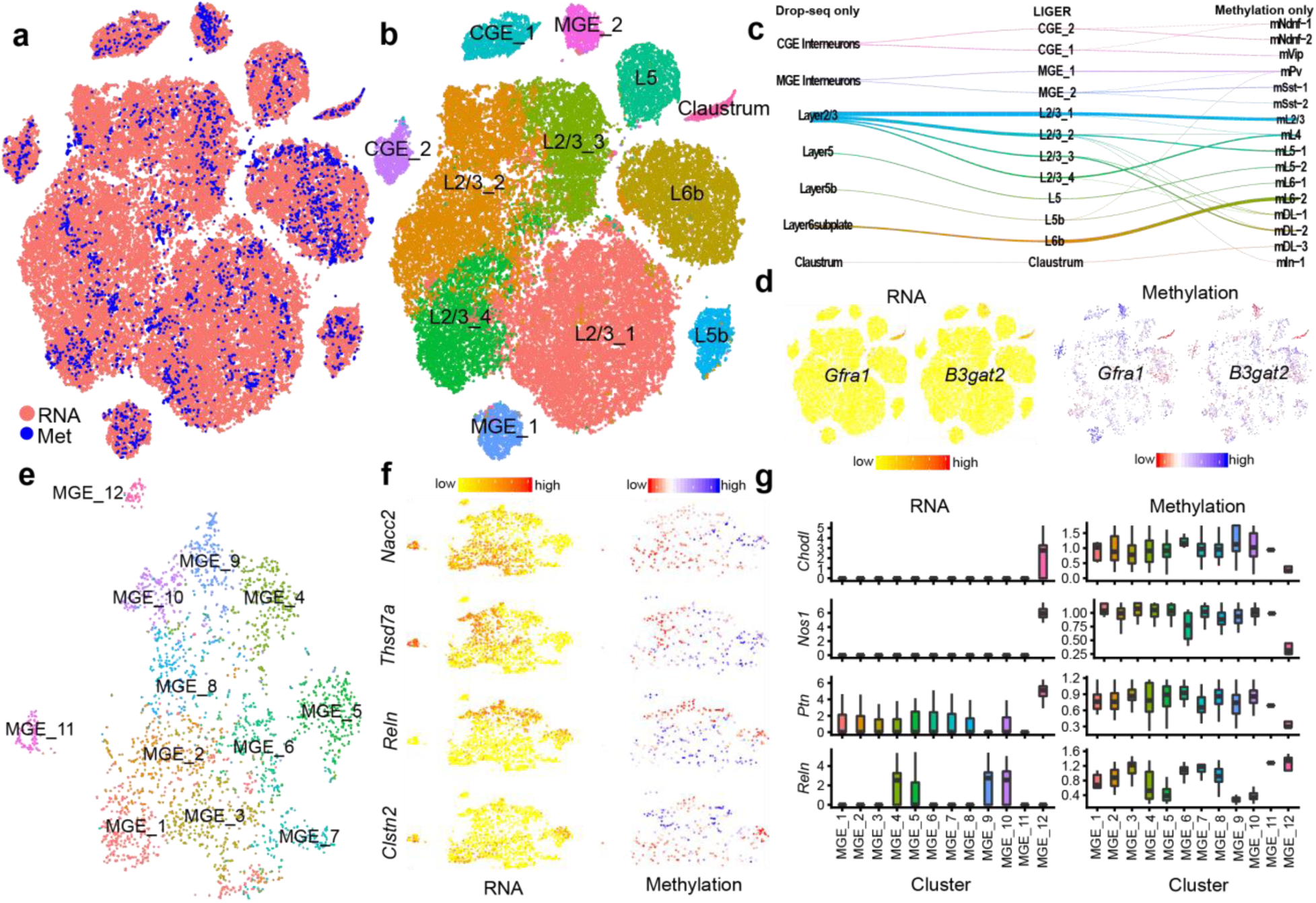
Defining cortical cell types using both scRNAseq and DNA methylation. **(a)-(b)** t-SNE visualization of LIGER analysis of single-cell RNAseq data (Saunders et al., 2018) and methylation data (Luo et al., 2017) from mouse frontal cortex, colored by modality **(a)** and LIGER cluster assignment **(b)**. **(c)** River plot showing relationship between published cluster assignments of RNA and methylation data and LIGER joint clusters. **(d)** Expression and methylation of two claustrum markers. **(e)** t-SNE representation of the LIGER subcluster analysis of MGE interneurons. **(f)** Expression and methylation of 4 marker genes for different MGE subpopulations. **(g)** Boxplots of expression and methylation markers for Sst-Chodl cells (cluster MGE_12).

Annotation of clusters defined solely by methylation data can be challenging, because most known markers of cell-type identity are mRNAs or proteins. Our joint analysis, utilizing small correlations across thousands of genes, allowed us to clarify the identities of some of the originally reported methylation clusters. First, we found that a cluster annotated as “deep layer cluster 3” aligned uniquely to an RNA-seq cluster that we previously had annotated as claustrum (Saunders et al., 2018) (**Figures 6C, D**). Second, a cluster annotated as “layer 6 cluster 1” aligned with a cluster that we identified as layer 5b. The canonical marker genes have relatively low overall methylation levels, making it challenging to assign the identity of this cell type from methylation alone. However, the expression of several specific layer 5b marker genes, most notably *Slc17a8* (Sorensen et al., 2015), and their corresponding low methylation pattern in the aligned cluster mL6-1 cells, enabled us to confirm this assignment (**Figure S5A**).

Consistent with previous large-scale, whole-tissue single-cell studies (Saunders et al., 2018), we found that a second level of clustering yielded additional substructure that was not visible in the global analysis. We performed four sub-analyses of the dominant cell types in the frontal cortex: CGE-derived interneurons, MGE-derived interneurons, superficial excitatory neurons, and deep-layer excitatory neurons (**Figures S5C-S5E**). From this second round of analysis, we identified a total of 37 clusters across the cortex. We found especially interesting substructure in the analysis of MGE-derived interneurons, where we identified 11 distinct populations, considerably more than was possible using the methylation data alone (**Figure 6E**). Examining expression and methylation of marker genes confirmed that these populations are real and not simply forced alignment (**Figure 6F**). The inclusion of the scRNA-seq dataset gave greater power to detect, within the methylation dataset, very rare cell types described by previous scRNA-seq studies of cortex. We were able to credibly identify 25 methylation profiles corresponding to an interneuron population defined by expression of *Pvalb* and *Th* (**Figure S5B**), as well as 5 profiles aligning to the cluster expressing *Sst* and *Chodl* (**Figure 6G**). The *Sst/Chodl* cells showed low methylation of *Nos1* and the very specific marker *Chodl*, and high methylation of *Reln*, consistent with this population’s previously established transcriptional phenotype (**Figure 6G**). Together, these results indicate that epigenetic and expression data produce highly concordant neural cell-type definitions, and even the finest distinctions among neural cell types defined from gene expression can be reflected by DNA methylation. Additionally, our increased sensitivity to detect methylation populations underscores the power of our integrative approach.

Our joint cluster definitions offer an opportunity to investigate the regulatory relationship between expression and methylation at cell-type-specific resolution. We first aggregated the gene expression and methylation values within each cluster, then calculated correlation between the expression of each gene and its gene body methylation levels across the set of clusters. Consistent with previous reports, we found an overall negative relationship between methylation and expression **(Figure 7A).** We observed a slight relationship with gene length: longer genes showed stronger negative correlation with gene expression (**Figure 7A**). This length relationship fits the pattern expected from a known mechanism of gene repression by DNA methylation, in which the MECP2 protein binds methylated nucleotides (Fasolino and Zhou, 2017). The degree of MECP2 repression has been shown to be proportional to the number of methylated nucleotides, which is strongly related to gene length (Kinde et al., 2016). However, the length of a gene also affects the amount of measured methylation signal, because long genes with more nucleotides are more likely to be captured in the sparse sampling of sequencing reads from a given cell, so we cannot completely rule out the influence of technical factors in this observed length relationship.

**Figure 7:**
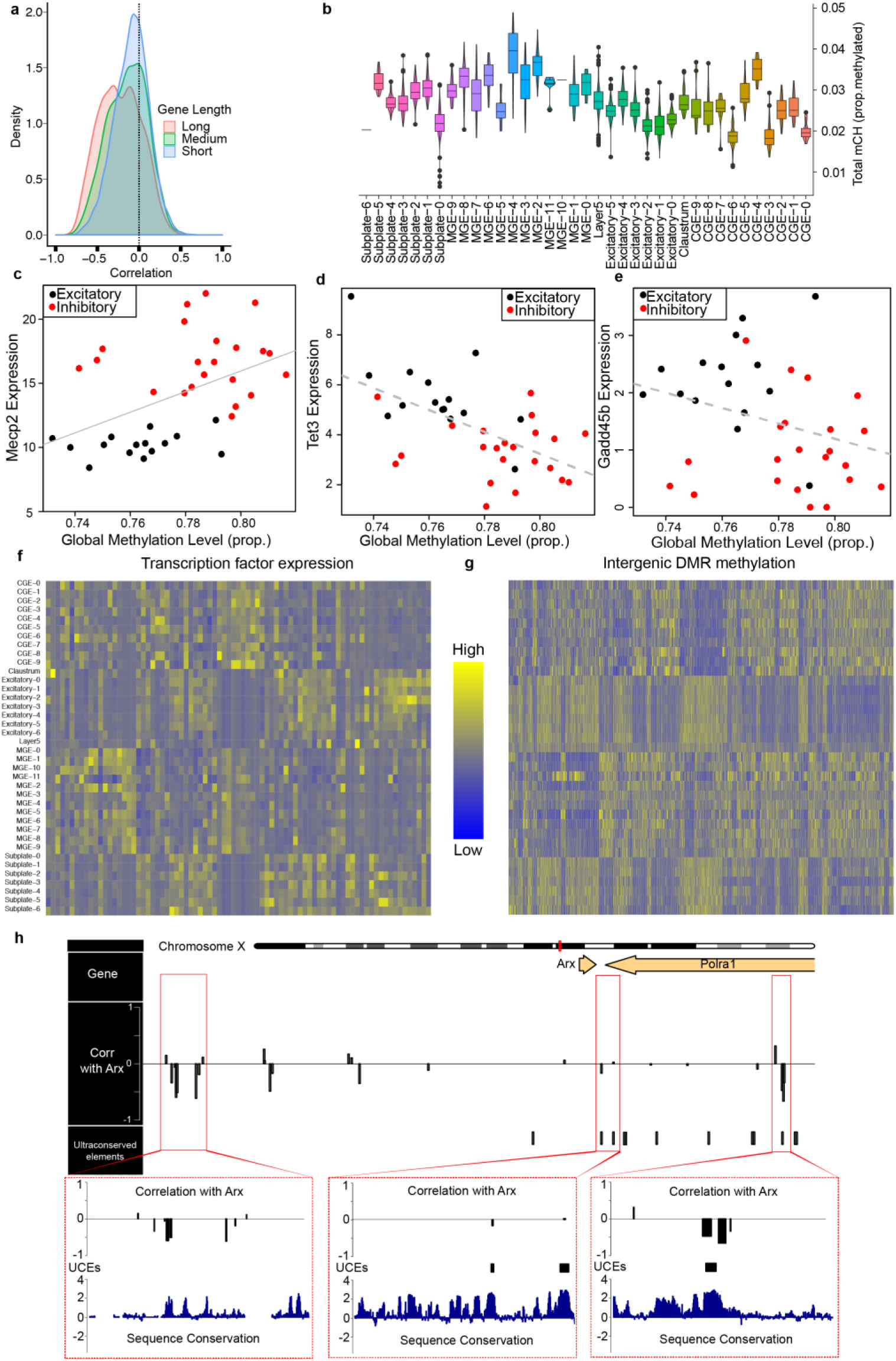
Investigating the connection between DNA methylation and gene expression. **(a)** Density plot of the correlation between gene body methylation and expression for short, medium, and long genes. **(b)** Violin plots showing the wide range of global methylation levels across neural cell types defined by the LIGER analysis in **Figure 6**. **(c)-(e)** Scatter plots of global methylation and aggregate expression for **(c)** *Mecp2*, **(d)** *Tet3*, and (**e**) *Dnmt3a* across our joint neural cell clusters. **(f)-(g)** Heatmaps of transcription factor expression **(f)** and methylation of intergenic regions predicted to be bound by these transcription factors **(g)**. **(h)** Genome browser view showing locations of differentially methylated regions near the *Arx* locus and their correlation with the expression of *Arx*. The bars indicate sign and magnitude of the correlation. The 3 bottom panels show zoomed-in views of three clusters of DMRs.

We observed a wide range of global methylation levels across our set of clusters, with inhibitory interneurons exhibiting the highest amount of methylation, as noted in the original analysis of the methylation data alone (Luo et al., 2017) (**Figure 7B**). Reasoning that this large dynamic range provides an opportunity to investigate the basic molecular machinery involved in regulating methylation, we correlated the expression of several key genes with the total amount of methylation per cell. We found that expression of *Mecp2* correlates strongly (ρ = 0.46, *p* = 0.0039) with total methylation level **(Figure 7C).** This pattern supports a model in which MECP2 represses gene expression by specifically binding to methylated nucleotides (Kinde et al., 2016), creating a stoichiometric requirement for increased *Mecp2* expression in cells with higher overall methylation levels. In addition, we found that *Tet3*, a gene involved in demethylation by conversion of 5mC to 5hmC, is strongly anticorrelated (ρ = −0.57, *p* = 0.0002) with global methylation level (**Figure 7D**). Intriguingly, the other TET genes do not show any anticorrelation with global methylation despite similar overall expression levels to *Tet3* (**Figure S6A** and **S6B**). This suggests that *Tet3* could be the dominant TET protein involved in actively regulating global methylation in mature neurons. *Gadd45b*, a gene with a well-established role in demethylating DNA in neurons (Bayraktar and Kreutz, 2018), also shows a strong negative relationship (ρ = −0.30, *p* = 0.0685) with total methylation. Consistent with our observation that both *Tet3* and *Gadd45b* are anticorrelated with total methylation, *Gadd45b* is thought to regulate DNA demethylation by recruiting TETs (Bayraktar and Kreutz, 2018). By contrast, none of the DNA methyltransferase enzymes (DNMTs) are strongly related to overall methylation level (**Figures S6C-S6E**). These analyses show the value of an integrated analysis to formulate hypotheses about the mechanisms by which expression and methylation are regulated.

Our integrated single-cell methylation and gene expression analysis could also enable the identification of intergenic regulatory elements that regulate cell-type-specific gene expression. Although previous studies have identified regulatory elements in uniform cell populations using bulk ChIP-seq, ATAC-seq, and bisulfite sequencing data (Consortium, 2012), our knowledge of cell-type-specific regulation in a complex tissue like brain is limited. Deciphering how epigenetic modification of specific sequence elements regulates gene expression is fundamental to understanding the establishment and maintenance of cell identity. In addition, locating enhancers and linking them to specific populations of brain cells is an important first step for unraveling the effects of genetic variants—many of which fall in intergenic regions—linked to neuropsychiatric disorders.

We identified a set of sequence elements that: (1) showed no overlap with any annotated genes or promoters; (2) showed significant cell-type-specific DNA methylation; (3) contain a conserved sequence element; (4) contain a binding motif for a transcription factor that is expressed in the dataset; and (5) have a methylation profile that is anticorrelated with the expression of the transcription factor whose binding motif occurs in the region (**Figure 7F; Methods**). **Figure 7G** shows the expression of the transcription factors and methylation of the top corresponding sequence elements for each TF. These intergenic elements represent strong candidates for cell-type-specific transcriptional regulatory elements, with available, unmethylated transcription factor binding motifs in the cell types where the corresponding transcription factors are most highly expressed.

Finally, our integrated definition of cell types from methylation and expression allowed us to examine the relationship between intergenic methylation and the expression of nearby genes. Identifying promoters or enhancers that drive expression of cell-type-specific genes is an essential part of designing adeno-associated viruses (AAVs) that target specific neuronal populations (Chan et al., 2017; Dimidschstein et al., 2016). We examined the expression of the *Arx* gene and methylation of neighboring loci across our joint clusters. Since *Arx* is a transcription factor that is specifically expressed in MGE-derived interneurons, identifying enhancers that regulate *Arx* could prove a useful tool for targeting this specific neuronal population. The *Arx* locus is remarkable in that 8 ultraconserved elements (UCEs)— long stretches of sequence showing perfect conservation among human, mouse, and rat (Bejerano et al., 2004; Colasante et al., 2008)—are located in the < 1 megabase region around the gene. Several distal regulatory elements, including some located within neighboring UCEs, have recently been demonstrated to regulate *Arx* expression (Colasante et al., 2008; Dickel et al., 2018). To nominate putative elements regulating *Arx*, we correlated *Arx* expression and methylation of nearby differentially methylated regions (DMRs) across our joint clusters (**Figure 7H**). We observed several clusters of DMRs whose methylation is anticorrelated with *Arx* expression, a pattern expected if hypo-methylation within certain cell types makes available a regulatory element that enhances *Arx* expression. One of these anticorrelated DMRs is a validated *Arx* enhancer (Dickel et al., 2018) that overlaps with an ultraconserved element just downstream of the end of the *Arx* gene (**Figure 7H**, middle). Another pair of DMRs strongly correlated with *Arx* expression overlap a UCE further downstream of *Arx* (**Figure 7H**, right). A third group of DMRs upstream of the *Arx* site lies in a region of very high conservation (though not a UCE), with three clear spikes in conservation that align precisely with the locations of the DMRs (**Figure 7H**, left). In summary, these DMRs represent strong candidates for putative elements regulating *Arx* expression, highlighting the value of our integrative approach for investigating gene regulatory mechanisms.

## Discussion

A credible definition of cell type requires distinguishing the invariant properties of cell identity from the dispensable across a myriad of settings and measurements. LIGER promises to be a broadly useful analytical tool for such efforts because of several key technical advantages. First, the nonnegativity constraint (i.e., metagene expression levels are never negative) yields interpretable factors, such that each factor generally corresponds to a biologically meaningful signal, like a collection of genes defining a particular cell type. The dataset-specific metagenes readily provide important biological insights into cell-type-specific differences across datasets, as we have shown in our analyses of sexual dimorphism in the bed nucleus of the mouse and in our comparative analysis of substantia nigra cell types across individual human postmortem donors. Second, the inclusion of dataset-specific terms allows us to identify dataset differences, rather than attempting to force highly divergent datasets into a completely shared latent space. This feature was especially useful in comparing and contrasting cell populations in the bed nucleus with related populations in the frontal cortex and striatum, as it allowed increased sensitivity to detect corresponding cell types from a small shared signal, while also reducing the inference of spurious connections between datasets. Finally, LIGER’s inference of both shared and dataset-specific factors enables a more transparent and nuanced definition of *how* cells correspond across datasets. In cases where complete correspondence is not necessarily expected—such as connecting fully differentiated cells to progenitors or relating pathological cells to healthy counterparts—a characterization of the metagenes that both unite and separate such populations is crucial.

We envision LIGER serving several important needs in neurobiology, beyond its capacity to better define cell types. First, a key opportunity in single-cell analysis is the identification of cell-type-specific gene expression patterns associated with disease risk, onset and progression in human tissue samples. Early efforts at such investigation have yielded some exciting results (Keren-Shaul et al., 2017), but increased discovery is likely possible with robust integrative analysis of many tissue donors. In addition to the identification of disease-relevant cell states, such analyses will also be indispensible to the localization of genetic risk loci for neuropsychiatric diseases to specific human cell types. Second, the integration of data from epigenomic and transcriptomic datasets provides a path towards nominating functional genomic elements important in cell-type-specific gene regulation. Such elements are compelling candidates for cell-type-specific enhancers to drive expression of genetic tools in specific subsets of brain cells. Their identification may also prove to be a valuable means of narrowing the search for causative alleles at specific genetic risk loci. Finally, as efforts to develop *in vitro* models of complex brain tissues continue to become more sophisticated (Birey et al., 2017; Quadrato et al., 2017), single-cell gene expression measurements, together with an integrative analysis like LIGER, will help provide systematic, information-rich comparisons of such models with their *in vivo* counterparts. To facilitate adoption of the tool in the community, we have developed <u>an R-package</u> that supports analysis of large-scale datasets and includes ancillary functions for tuning algorithmic parameters, visualizing results, and quantifying integrative performance. We hope its widespread deployment opens many exciting new avenues in single-cell biology.

## Author Contributions

J.W. and E.M. designed the study and wrote the paper. J.W. derived and implemented the computational algorithms, with help from V.K. J.W., V.K., and E.M. performed the analyses. J.W. and V.K. wrote the R package. A.F., C.M., and C.V. performed the dissections, nuclear isolations, and library preparations from mouse bed nucleus and human substantia nigra.

## Acknowledgments

We thank Aleks Goeva for advice on computational algorithms. This work was supported by the Stanley Center for Psychiatric Research, the Chan Zuckerberg Initiative (grant numbers 2017-175259 and 2018-183155), and NIH/NIMH BRAIN Grant 1U19MH114821.

## Methods

### LIGER Workflow

A typical workflow for Integrating multiple single-cell datasets using the LIGER software package consists of the following steps:

- Dataset preprocessing to produce a raw digital gene expression (DGE) matrix.
- Variable gene selection (Saunders et al., 2018), normalization by number of UMIs, and scaling of individual genes. We scale but do not center gene expression because NMF requires non-negative values.
- Identifying shared and dataset-specific factors through integrative non-negative matrix factorization (iNMF).
- Jointly clustering cells and normalizing factor loadings.
- Visualization using t-SNE or UMAP and analysis of shared and dataset-specific marker genes.

The important components of each step are described in greater detail below, while vignettes which describe specific commands for analyses included in **Figure 2** are available online at https://macoskolab.github.io/liger/.

### Performing Integrative Nonnegative Matrix Factorization Using Block Coordinate Descent

We developed a novel block coordinate descent algorithm for performing integrative non-negative matrix factorization (Yang and Michailidis, 2016). This approach learns a set of latent metagene factors, each with both shared and dataset-specific components, to approximate the original datasets. To estimate these matrix factors, we minimize the following objective:

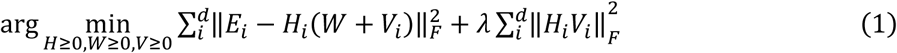

This approach attempts to reconstruct each of the original datasets *E_i_* (of dimension *n_i_* × *m*) using lower dimensional matrices *H_i_*, *W*, and *V_i_*, such that *Ei* ≈ *H_i_*(*W* + *V_i_*), where all factor matrices are constrained to be non-negative. Note that *W* is shared across all datasets *i* = 1… *d*, while the *H_i_* and *V_i_* matrices are unique to each dataset. The inner dimension *k* of these factors can be interpreted as the number of “metagenes” (or conversely “metacells”) used to represent the datasets.

We divide the variables into 2*d* + 1 blocks (corresponding to *H* and *V* for each dataset, as well as *W*) and perform block coordinate descent, iteratively minimizing the objective with respect to each block, holding the others fixed. We iterate:

1. 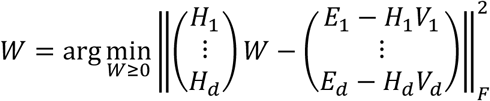
2. 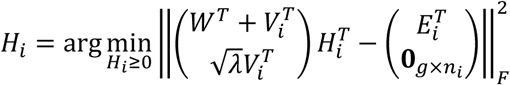
3. 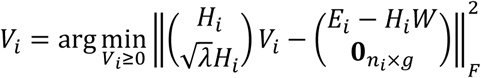 (**0**_*c×d*_ is the zero matrix of dimension *c* × *d*).

until convergence. Each of the optimization subproblems above requires solving a nonnegative least squares problem; we use the fast block principal pivoting algorithm developed by Kim et al. (Kim, 2014) to solve each of these subproblems exactly. As described in Kim et al, our block coordinate descent algorithm satisfies the requirements of the theorem of Bertsekas (Bertsekas, 1999), because each of the subproblems is convex with respect to the block of variables being optimized. Thus, the algorithm is guaranteed to converge to a fixed point (local minimum). In contrast, the multiplicative updates often used for NMF-like optimization problems do not have a convergence guarantee. Additionally, because we solve each subproblem exactly at each iteration, the algorithm converges very quickly; previous empirical benchmarks (Kim, 2014) and our own have shown that block coordinate descent algorithms for NMF generally converge in many fewer iterations than multiplicative updates (**Figure S1A-B**). We developed an efficient implementation of the algorithm using the Rcpp package in R.

### Efficient Updating for New K, New Data, and New λ

We adapted a method, previously developed for regular NMF (Kim, 2014), for rapidly updating a factorization given new data or new values of *k* or λ. Suppose we have optimized (1) with *k*_1_ factors to give matrices 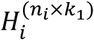, 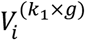, and *W*^(k_1_×*g*)^. To efficiently compute a factorization with *k*_2_ factors, we consider two cases: *k*_2_ > *k*_1_ and *k*_2_ < *k*_1_. If *k*_2_ > *k*_1_, we initialize the *k*_2_ − *k*_1_ new factors to factorize the residual from the previous solution, then solve using alternating nonnegative least squares as before. For *k*_2_ > *k*_1_, we pick the *k*_2_ factors that make the largest contributions to the factorization and solve as before. The following algorithm formalizes this approach.

#### Algorithm 1: Updating the factorization with a new *k*

1. Initialize *W^new^*, 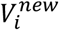, 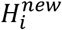 with random nonnegative values.
2. If *k*_2_ > *k*_1_:

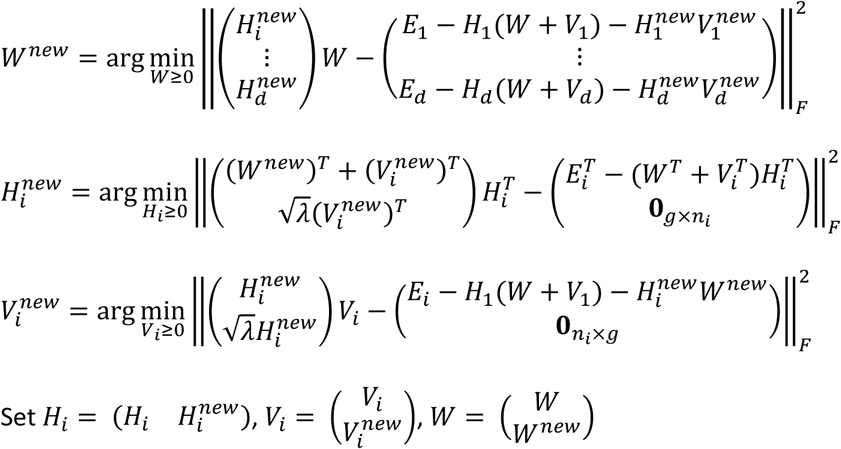
3. If *k*_2_ < *k*_1_: Let 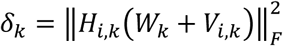. Choose the factors with the largest *k*_2_ values of δ_*k*_. Call the corresponding elements of *W*, *H_i_*, and *V_i_* Set 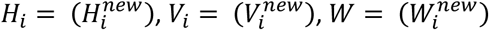
4. Perform block coordinate descent optimization using alternating nonnegative least squares until convergence.

Similarly, we can efficiently compute a factorization with a new λ value. Assume a previous optimization with *k* factors and λ_1_, and with matrices 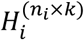, 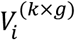, and *W*(*k×*). To compute a new factorization with λ_2_ > λ_1_, we can use the matrices *H_i_* and *W* and simply reoptimize with the new λ value. Empirical benchmarks for updating factorizations with new *k* and λ values are shown in Figure S1C-D.

Suppose we have optimized (1) with *k* factors to give matrices 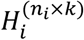, 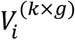, and *W*^(*k×g*)^. To efficiently compute a factorization incorporating new data from the same condition as an existing dataset, we use the previous *W* and *V* matrices as initial values and find the optimal *H* given this starting point. Given a new dataset altogether, we initialize the *W* and *V* matrices using the values from the dataset that we expect *a priori* to be the most similar and find the optimal *H* given these values. To re-run on a subset of the data, we use the *W* and *V* matrices as a starting point and simply drop the *H* rows corresponding to the omitted data. For each case, we subsequently perform optimization using the same block coordinate descent strategy as described above. Algorithm 2 summarizes this approach. Empirical benchmarks for updating factorizations with subsets of data and new data are shown in Figure S1E-F.

#### Algorithm 2: Adding new data to the factorization

1. Given new data from one of the datasets already factorized or a completely new dataset, initialize *W* and *V_i_* to their previously found values (if a completely new dataset, choose *V_i_* from the dataset that is expected to be most similar), then solve:

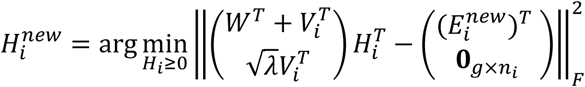 Set 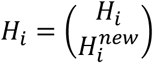.
2. To update the factorization for only a subset of the data, initialize *H_i_* by simply dropping the *H_i_* rows corresponding to the omitted data.
3. Perform block coordinate descent optimization using alternating nonnegative least squares until convergence.

### Heuristics to guide choice of *k* and λ

We devised novel heuristics to aid in selecting the number of factors *k* and the tuning parameter λ. To choose *k*, we calculate the Kullback-Leibler divergence (compared to a uniform distribution) of the factor loadings for each cell and plot the median across cells as a function of *k*. We then try to observe a saturation point in the curve, corresponding to the point at which more factors do not significantly change the sparsity of the factor loadings or correspondingly increase the median KL divergence. The intuition behind this heuristic is that, if the number of factors is too low, factors will encode combinations of clusters, and thus cells will load on multiple factors, with the distribution of factor loadings for a particular cell approaching a uniform distribution. As the number of factors approaches the “true” number of clusters in the dataset, each cell will generally be expected to load on only a few factors. To select an appropriate lambda, we plot the alignment metric (after performing factorization and basic alignment with default parameters) as a function of lambda and again choose the minimum value at which the alignment metric saturates. KL divergence curves and alignment-lambda curves for two benchmark datasets are showing in **Figures S1G-H**.

### Shared factor neighborhood clustering and factor normalization

After optimizing the iNMF objective function, we use the factor space to identify corresponding cell types across datasets (**Figure 1C**). Because of the parts-based nature of the factorization, cells can be clustered by simply assigning each cell to the factors on which they have the highest loading. This is a common way of performing clustering using NMF representations (Xu, 2003). If we perform such simple assignment, we get clusters that correspond across datasets, because the factors represent the same signal in each dataset. However, we noticed that simple maximum factor assignments sometimes produced spurious alignments in highly divergent datasets. Therefore, we developed a novel clustering strategy that better leverages the dataset-specific information in the factorization to increase the robustness of the joint clustering results. We build a *shared factor neighborhood* graph in which we connect cells across datasets that have similar factor loading patterns, then identify joint clusters by performing community detection on this graph.

More specifically, we build the shared factor neighborhood (SFN) graph as follows:

1. Build *k*-nearest neighbor graphs separately for each dataset using the *H* factor loadings
2. Annotate each cell *i* with the *H* factor on which the cell has the highest loading; call this *F*(*i*). We scale each factor to unit variance before assigning *F*(*i*) values, as is standard with NMF (Xu, 2003).
3. For each cell *i*, collect the “factor neighborhood” vector *FN*(*i*) by computing a histogram of *F*(*i*) for each of its *k* nearest neighbors.
4. Calculate Manhattan distance between pairs of cells (*i, j*) across (and within) datasets using the factor neighborhood vectors *FN*(*i*) and *FN*(*j*)
5. Connect pairs of cells with low distance; these represent cells with shared factor neighborhoods.

We then perform Louvain community detection on the graph to jointly identify cell clusters across datasets.

One intuition behind this construction is that, *within* an individual dataset, it is much more likely for an individual cell to have a spurious maximum factor loading than for an entire neighborhood of cells to be incorrectly assigned. Thus, leveraging the neighborhood of a cell in assigning it to a cluster increases the robustness of the assignment. This approach is synergistic with iNMF, which reconstructs each dataset separately, accurately preserving the structure of individual datasets. A recent paper used a related idea to refine cluster assignments from NMF by taking into account the neighborhood of each data point (Tripodi, 2016). Additionally, our graph construction greatly reduces the chances of spurious matches *across* datasets, because even if a cell type spuriously loads on the same factor as a different cell type in another dataset, they are unlikely to have the same factor neighborhoods.

After performing SFN clustering, we choose a reference dataset (by default the dataset with the largest number of cells) and normalize the quantiles of the factor loadings for each joint cluster in the other datasets to match the quantiles of the reference dataset for that joint cluster. We require that each dataset have a minimum number of cells assigned to a particular cluster (default: 2 cells); cells not satisfying this requirement are not normalized.

### Calculating alignment and agreement metrics

We calculated the alignment metric as defined in Butler et al. (Butler et al., 2018). To quantify how well the integrated factor space respects the geometry of each individual dataset, we calculate an “agreement metric” as follows. We first apply a dimensionality reduction technique to each dataset separately. Using these low-dimensional representations, we build a *k*-nearest neighbor graph for each dataset. We also build *k*-nearest neighbor graphs for each dataset using the joint, integrated space—the normalized *H* factor loadings from LIGER or the aligned canonical components from Seurat. We then count how many of each cell’s nearest neighbors in the graphs built from the separate low-dimensional representations are also nearest neighbors in the graphs built from the integrated low-dimensional representations. We found that PCA and NMF produced significantly different graphs for different joint dimensionality reduction and alignment techniques, so to ensure a fair comparison, we compared the graphs built from the aligned Seurat space (CCA-based) with graphs built from PCA, and compared the graphs built from the aligned LIGER factor loadings (iNMF-based) with NMF.

For comparisons with Seurat, we followed all documented procedures for running the method, including choosing the number of canonical components. For the PBMC and pancreas datasets, we reproduced the analyses as described by Butler et al. and subsequent Seurat tutorials (Butler et al., 2018).

### Analysis of bed nucleus data

We performed a first round of LIGER analysis on BNST nuclei to identify neurons, then restricted subsequent analyses to neuronal cells. We next assigned neuronal populations to anatomical locations using in situ staining data from the Allen Brain Atlas (**Figure S3**), then performed a second round of clustering on neurons we could assign to the bed nucleus. Clustering of bed nucleus neurons identified 32 populations, four of which were removed because of doublet signatures. We found that it was not necessary to quantile normalize the factor spaces for the bed nucleus analyses, as iNMF alone provides sufficient batch effect correction in this case. We used the iNMF factorization to identify genes with dimorphic expression by simply taking the genes with nonzero loadings on the dataset-specific factors *V*. As an additional filtering step, for gene *i* loading on dataset-specific factor *j*, we removed *i* if it had a log fold change of less than 0.75 or fewer than 30 UMIs in the cells with their highest factor loading values on factor *j*. Noting that each factor generally loads on only a single cluster, we then assigned dimorphic genes to the cluster on which the corresponding factor had the highest average loading. To rank dimorphic genes by their degree of dimorphism (as shown in **Figure 3K**), we calculated the difference, for each gene, between male and female dataset specific factor loadings for the factors on which the gene is dimorphic.

For the comparison of BNST_Vip nuclei with CGE-derived cortical interneurons, we randomly sampled 50 cells from each of the 11 published subclusters of frontal cortex global cluster 1 (Saunders et al., 2018), to maintain cell-type diversity in the downsampled set. These 550 CGE neurons were analyzed with the 139 BNST_Vip profiles with LIGER according to parameters in Table S1. For the comparison of the *Ppp1r1b+* BNST populations with striatal SPNs, we sampled 2000 dSPNs (published global cluster 10), 2000 iSPNs (published global cluster 11), and 1,084 eSPNs (subclusters 1-5 of global cluster 13). These 5,084 SPNs were analyzed with the 3,652 *Ppp1r1b+* nuclei from the BNST analysis using LIGER with the parameters in **Table S1**.

### Analysis of substantia nigra data

To analyze the human substantia nigra data across donors, we used a separate dataset-specific factor matrix for each human donor. We performed two rounds of clustering with LIGER, first identifying the main cell classes (neurons, endothelial cells, astrocytes, oligodendrocytes, and microglia), then clustering each cell type again to identify additional substructure within these classes (Saunders et al., 2018). For the cross-species analysis, we determined homology relationships using Jackson Laboratories annotations (http://www.informatics.jax.org) and included only genes with one-to-one homologs. We used LIGER to integrate each broad cell class separately, and used two dataset-specific factor matrices (one for each species), rather than treating the data from each human donor as a separate dataset. For the cross-species analysis, we performed variable gene selection on each species separately, then took the union of variable genes across both species.

For the identification of gene ontology categories with conserved patterns of expression across species, we used GOrilla (Eden et al., 2009) in ranked gene list mode, using default settings, to perform gene ontology enrichment analysis and ReviGO (Supek et al., 2011) to summarize and visualize the results.

### Analysis of STARmap data

We downloaded the published STARmap expression data and segmented cell boundaries, then normalized and scaled the STARmap gene counts in the same manner as the Drop-seq data of mouse frontal cortex (Saunders et al., 2018). Because the STARmap data assays pre-selected markers, we did not perform variable gene selection, but used all genes that were present in both the Drop-Seq and STARmap data. We used the entire frontal cortex dataset from a previously published Drop-Seq atlas of mouse brain. We performed two levels of analysis using LIGER, first jointly identifying broad cell classes, then performing a second round of LIGER analysis on excitatory neurons, inhibitory neurons, and glia.

### Analysis of methylation data

We downloaded the publicly available gene-level mCH fractions from 3378 frontal cortex profiles (Luo et al., 2017). Because gene expression and gene body mCH are generally anticorrelated, we created gene-level methylation features that correspond to gene expression features by calculating

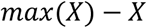

where *X* is the matrix of mCH values. Because the data from Luo et al. contains only neuronal cells, we used only cells annotated as neurons from the Saunders et al. frontal cortex Drop-seq dataset.

Because the distribution of methylation values is very different from scRNA-seq data, we selected genes by performing a Kruskal-Wallis test on methylation and RNA data separately using the published clusters, then taking the intersection of the top 8000 RNA and methylation markers. We found that, since the methylation data are not sparse (i.e., very few values are exactly 0), the NMF factor loadings need to be centered as well as scaled during the SFN clustering procedure. We also noted that the NMF algorithm converges extremely slowly compared to our ALS implementation when running on the non-sparse methylation data. We used the methylpy Python package to calculate methylation levels for the set of differentially methylated regions identified in the original analysis of the methylation data (Luo et al., 2017). We performed an analysis of transcription factor binding using FIMO (Grant et al., 2011) with default settings and the mouse transcription factor binding motifs from the latest version of the non-redundant JASPAR database (Khan et al., 2018). We annotated the DMRs according to the nearest gene using the *annotate* R package (https://www.bioconductor.org/packages/release/bioc/html/annotate.html)

### Generation of BNST nuclei profiles

All experiments were approved by and in accordance with Broad Institute IACUC protocol number 0120-09-16. Mouse brains (four male replicates, two female replicates) were perfused and flash frozen in liquid nitrogen, and mounted on a cryostat. Tissue was sliced from the anterior until reaching the target region of interest (**Figure S3A**). To dissect a tissue segment, the block face was punched with a 1 mm biopsy punch, and a 400-micron slice was made to liberate the circular tissue segments from the remaining brain. Nuclei were extracted from frozen tissue according to a protocol generously provided by the McCarroll lab (http://mccarrolllab.com/no-access/protocols/), and loaded into the 10x Chromium V2 system. Reverse transcription and library generation were performed according to the manufacturer’s protocol.

### Generation of profiles from human substantia nigra

Substantia nigra (SN) tissue from seven de-identified human donors was obtained from the University of Maryland Brain Bank, through the NIH Neurobank (exempted from human subjects research by ORSP, project number NHSR-4235). Although all seven donors were coded as neurotypical controls, information provided with the tissue revealed that donor 5828 suffered a traumatic brain injury at the time of death, while donor 5840 was diagnosed at autopsy with cerebral amyloid angiopathy.

Frozen tissue samples were sectioned on a cryostat to visualize pigmentation of the pars compacta, and manually dissected to remove as much surrounding tissue as possible. Nuclei from the dissected tissue segments were then immediately isolated (Saunders et al., 2018) and profiled on the 10x Chromium V2 system, according to the manufacturer’s protocol.

## References

Allen, L.S., and Gorski, R.A. (1990). Sex difference in the bed nucleus of the stria terminalis of the human brain. The Journal of comparative neurology 302, 697-706.

Amir, S., Lamont, E.W., Robinson, B., and Stewart, J. (2004). A circadian rhythm in the expression of PERIOD2 protein reveals a novel SCN-controlled oscillator in the oval nucleus of the bed nucleus of the stria terminalis. The Journal of neuroscience : the official journal of the Society for Neuroscience 24, 781-790.

Arendt, D., Musser, J.M., Baker, C.V.H., Bergman, A., Cepko, C., Erwin, D.H., Pavlicev, M., Schlosser, G., Widder, S., Laubichler, M.D., et al. (2016). The origin and evolution of cell types. Nat Rev Genet 17, 744-757.

Baron, M., Veres, A., Wolock, S.L., Faust, A.L., Gaujoux, R., Vetere, A., Ryu, J.H., Wagner, B.K., Shen-Orr, S.S., Klein, A.M., et al. (2016). A Single-Cell Transcriptomic Map of the Human and Mouse Pancreas Reveals Inter- and Intra-cell Population Structure. Cell Syst 3, 346-360 e344.

Bayless, D.W., and Shah, N.M. (2016). Genetic dissection of neural circuits underlying sexually dimorphic social behaviours. Philos Trans R Soc Lond B Biol Sci 371, 20150109.

Bayraktar, G., and Kreutz, M.R. (2018). The Role of Activity-Dependent DNA Demethylation in the Adult Brain and in Neurological Disorders. Front Mol Neurosci 11, 169.

Bejerano, G., Pheasant, M., Makunin, I., Stephen, S., Kent, W.J., Mattick, J.S., and Haussler, D. (2004). Ultraconserved elements in the human genome. Science 304, 1321-1325.

Bertsekas, D. (1999). Nonlinear Programming (Athena Scientific).

Biffi, A., and Greenberg, S.M. (2011). Cerebral amyloid angiopathy: a systematic review. J Clin Neurol 7, 1-9.

Birey, F., Andersen, J., Makinson, C.D., Islam, S., Wei, W., Huber, N., Fan, H.C., Metzler, K.R.C., Panagiotakos, G., Thom, N., et al. (2017). Assembly of functionally integrated human forebrain spheroids. Nature 545, 54-59.

Bruinsma, I.B., de Jager, M., Carrano, A., Versleijen, A.A., Veerhuis, R., Boelens, W., Rozemuller, A.J., de Waal, R.M., and Verbeek, M.M. (2011). Small heat shock proteins induce a cerebral inflammatory reaction. The Journal of neuroscience : the official journal of the Society for Neuroscience 31, 11992-12000.

Butler, A., Hoffman, P., Smibert, P., Papalexi, E., and Satija, R. (2018). Integrating single-cell transcriptomic data across different conditions, technologies, and species. Nat Biotechnol 36, 411-420.

Chan, K.Y., Jang, M.J., Yoo, B.B., Greenbaum, A., Ravi, N., Wu, W.L., Sanchez-Guardado, L., Lois, C., Mazmanian, S.K., Deverman, B.E., et al. (2017). Engineered AAVs for efficient noninvasive gene delivery to the central and peripheral nervous systems. Nat Neurosci 20, 1172-1179.

Colasante, G., Collombat, P., Raimondi, V., Bonanomi, D., Ferrai, C., Maira, M., Yoshikawa, K., Mansouri, A., Valtorta, F., Rubenstein, J.L., et al. (2008). Arx is a direct target of Dlx2 and thereby contributes to the tangential migration of GABAergic interneurons. J Neurosci 28, 10674-10686.

Consortium, E.P. (2012). An integrated encyclopedia of DNA elements in the human genome. Nature 489, 57-74.

Coskun, A.F., and Cai, L. (2016). Dense transcript profiling in single cells by image correlation decoding. Nature methods 13, 657-660.

Cusanovich, D.A., Hill, A.J., Aghamirzaie, D., Daza, R.M., Pliner, H.A., Berletch, J.B., Filippova, G.N., Huang, X., Christiansen, L., DeWitt, W.S., et al. (2018). A Single-Cell Atlas of In Vivo Mammalian Chromatin Accessibility. Cell 174, 1309-1324 e1318.

Dickel, D.E., Ypsilanti, A.R., Pla, R., Zhu, Y., Barozzi, I., Mannion, B.J., Khin, Y.S., Fukuda-Yuzawa, Y., Plajzer-Frick, I., Pickle, C.S., et al. (2018). Ultraconserved Enhancers Are Required for Normal Development. Cell 172, 491-499 e415.

Dimidschstein, J., Chen, Q., Tremblay, R., Rogers, S.L., Saldi, G.A., Guo, L., Xu, Q., Liu, R., Lu, C., Chu, J., et al. (2016). A viral strategy for targeting and manipulating interneurons across vertebrate species. Nature neuroscience 19, 1743-1749.

Dimou, L., Simon, C., Kirchhoff, F., Takebayashi, H., and Gotz, M. (2008). Progeny of Olig2-expressing progenitors in the gray and white matter of the adult mouse cerebral cortex. The Journal of neuroscience : the official journal of the Society for Neuroscience 28, 10434-10442.

Ding, S.L., Royall, J.J., Sunkin, S.M., Ng, L., Facer, B.A., Lesnar, P., Guillozet-Bongaarts, A., McMurray, B., Szafer, A., Dolbeare, T.A., et al. (2016). Comprehensive cellular-resolution atlas of the adult human brain. The Journal of comparative neurology 524, 3127-3481.

Dolatshad, H., Campbell, E.A., O'Hara, L., Maywood, E.S., Hastings, M.H., and Johnson, M.H. (2006). Developmental and reproductive performance in circadian mutant mice. Hum Reprod 21, 68-79.

Dong, H.W., and Swanson, L.W. (2004). Projections from bed nuclei of the stria terminalis, posterior division: implications for cerebral hemisphere regulation of defensive and reproductive behaviors. The Journal of comparative neurology 471, 396-433.

Eden, E., Navon, R., Steinfeld, I., Lipson, D., and Yakhini, Z. (2009). GOrilla: a tool for discovery and visualization of enriched GO terms in ranked gene lists. BMC Bioinformatics 10, 48.

Fasolino, M., and Zhou, Z. (2017). The Crucial Role of DNA Methylation and MeCP2 in Neuronal Function. Genes (Basel) 8.

Gierahn, T.M., Wadsworth, M.H., 2nd, Hughes, T.K., Bryson, B.D., Butler, A., Satija, R., Fortune, S., Love, J.C., and Shalek, A.K. (2017). Seq-Well: portable, low-cost RNA sequencing of single cells at high throughput. Nat Methods 14, 395-398.

Grant, C.E., Bailey, T.L., and Noble, W.S. (2011). FIMO: scanning for occurrences of a given motif. Bioinformatics 27, 1017-1018.

Gustafson, E.L., and Greengard, P. (1990). Localization of DARPP-32 immunoreactive neurons in the bed nucleus of the stria terminalis and central nucleus of the amygdala: co-distribution with axons containing tyrosine hydroxylase, vasoactive intestinal polypeptide, and calcitonin gene-related peptide. Exp Brain Res 79, 447-458.

Habib, N., Avraham-Davidi, I., Basu, A., Burks, T., Shekhar, K., Hofree, M., Choudhury, S.R., Aguet, F., Gelfand, E., Ardlie, K., et al. (2017). Massively parallel single-nucleus RNA-seq with DroNc-seq. Nat Methods.

Haghverdi, L., Lun, A.T.L., Morgan, M.D., and Marioni, J.C. (2018). Batch effects in single-cell RNA-sequencing data are corrected by matching mutual nearest neighbors. Nat Biotechnol 36, 421-427.

Hines, M., Allen, L.S., and Gorski, R.A. (1992). Sex differences in subregions of the medial nucleus of the amygdala and the bed nucleus of the stria terminalis of the rat. Brain Res 579, 321-326.

Hodge, R.D., Bakken, T.E., Miller, J.A., Smith, K.A., Barkan, E.R., Graybuck, L.T., Close, J.L., Long, B., Penn, O., Yao, Z., et al. (2018). Conserved cell types with divergent features between human and mouse cortex. bioRxiv.

Hrvatin, S., Hochbaum, D.R., Nagy, M.A., Cicconet, M., Robertson, K., Cheadle, L., Zilionis, R., Ratner, A., Borges-Monroy, R., Klein, A.M., et al. (2018). Single-cell analysis of experience-dependent transcriptomic states in the mouse visual cortex. Nat Neurosci 21, 120-129.

Johnson, W.E., Li, C., and Rabinovic, A. (2007). Adjusting batch effects in microarray expression data using empirical Bayes methods. Biostatistics 8, 118-127.

Keren-Shaul, H., Spinrad, A., Weiner, A., Matcovitch-Natan, O., Dvir-Szternfeld, R., Ulland, T.K., David, E., Baruch, K., Lara-Astaiso, D., Toth, B., et al. (2017). A Unique Microglia Type Associated with Restricting Development of Alzheimer's Disease. Cell 169, 1276-1290 e1217.

Khan, A., Fornes, O., Stigliani, A., Gheorghe, M., Castro-Mondragon, J.A., van der Lee, R., Bessy, A., Cheneby, J., Kulkarni, S.R., Tan, G., et al. (2018). JASPAR 2018: update of the open-access database of transcription factor binding profiles and its web framework. Nucleic Acids Res 46, D260-D266.

Kim, J.H., Yunlong; Park, Haesun (2014). Algorithms for nonnegative matrix and tensor factorizations: a unified view based on block coordinate descent framework. Journal of Global Optimization 58, 285-319.

Kim, S.Y., Adhikari, A., Lee, S.Y., Marshel, J.H., Kim, C.K., Mallory, C.S., Lo, M., Pak, S., Mattis, J., Lim, B.K., et al. (2013). Diverging neural pathways assemble a behavioural state from separable features in anxiety. Nature 496, 219-223.

Kinde, B., Wu, D.Y., Greenberg, M.E., and Gabel, H.W. (2016). DNA methylation in the gene body influences MeCP2-mediated gene repression. Proc Natl Acad Sci U S A 113, 15114-15119.

Klein, A.M., Mazutis, L., Akartuna, I., Tallapragada, N., Veres, A., Li, V., Peshkin, L., Weitz, D.A., and Kirschner, M.W. (2015). Droplet barcoding for single-cell transcriptomics applied to embryonic stem cells. Cell 161, 1187-1201.

Kudo, T., Uchigashima, M., Miyazaki, T., Konno, K., Yamasaki, M., Yanagawa, Y., Minami, M., and Watanabe, M. (2012). Three types of neurochemical projection from the bed nucleus of the stria terminalis to the ventral tegmental area in adult mice. The Journal of neuroscience : the official journal of the Society for Neuroscience 32, 18035-18046.

Lacar, B., Linker, S.B., Jaeger, B.N., Krishnaswami, S., Barron, J., Kelder, M., Parylak, S., Paquola, A., Venepally, P., Novotny, M., et al. (2016). Nuclear RNA-seq of single neurons reveals molecular signatures of activation. Nat Commun 7, 11022.

Lake, B.B., Ai, R., Kaeser, G.E., Salathia, N.S., Yung, Y.C., Liu, R., Wildberg, A., Gao, D., Fung, H.L., Chen, S., et al. (2016). Neuronal subtypes and diversity revealed by single-nucleus RNA sequencing of the human brain. Science 352, 1586-1590.

Lein, E.S., Hawrylycz, M.J., Ao, N., Ayres, M., Bensinger, A., Bernard, A., Boe, A.F., Boguski, M.S., Brockway, K.S., Byrnes, E.J., et al. (2007). Genome-wide atlas of gene expression in the adult mouse brain. Nature 445, 168-176.

Luo, C., Keown, C.L., Kurihara, L., Zhou, J., He, Y., Li, J., Castanon, R., Lucero, J., Nery, J.R., Sandoval, J.P., et al. (2017). Single-cell methylomes identify neuronal subtypes and regulatory elements in mammalian cortex. Science 357, 600-604.

Luo, J., Elwood, F., Britschgi, M., Villeda, S., Zhang, H., Ding, Z., Zhu, L., Alabsi, H., Getachew, R., Narasimhan, R., et al. (2013). Colony-stimulating factor 1 receptor (CSF1R) signaling in injured neurons facilitates protection and survival. J Exp Med 210, 157-172.

Macosko, E.Z., Basu, A., Satija, R., Nemesh, J., Shekhar, K., Goldman, M., Tirosh, I., Bialas, A.R., Kamitaki, N., Martersteck, E.M., et al. (2015). Highly Parallel Genome-wide Expression Profiling of Individual Cells Using Nanoliter Droplets. Cell 161, 1202-1214.

Manko, M., Bienvenu, T.C., Dalezios, Y., and Capogna, M. (2012). Neurogliaform cells of amygdala: a source of slow phasic inhibition in the basolateral complex. J Physiol 590, 5611-5627.

Mo, A., Mukamel, E.A., Davis, F.P., Luo, C., Henry, G.L., Picard, S., Urich, M.A., Nery, J.R., Sejnowski, T.J., Lister, R., et al. (2015). Epigenomic Signatures of Neuronal Diversity in the Mammalian Brain. Neuron 86, 1369-1384.

Moffitt, J.R., and Zhuang, X. (2016). RNA Imaging with Multiplexed Error-Robust Fluorescence In Situ Hybridization (MERFISH). Methods Enzymol 572, 1-49.

Mulqueen, R.M., Pokholok, D., Norberg, S.J., Torkenczy, K.A., Fields, A.J., Sun, D., Sinnamon, J.R., Shendure, J., Trapnell, C., O'Roak, B.J., et al. (2018). Highly scalable generation of DNA methylation profiles in single cells. Nat Biotechnol 36, 428-431.

Nery, S., Fishell, G., and Corbin, J.G. (2002). The caudal ganglionic eminence is a source of distinct cortical and subcortical cell populations. Nature neuroscience 5, 1279-1287.

Quadrato, G., Nguyen, T., Macosko, E.Z., Sherwood, J.L., Min Yang, S., Berger, D.R., Maria, N., Scholvin, J., Goldman, M., Kinney, J.P., et al. (2017). Cell diversity and network dynamics in photosensitive human brain organoids. Nature 545, 48-53.

Risso, D., Ngai, J., Speed, T.P., and Dudoit, S. (2014). Normalization of RNA-seq data using factor analysis of control genes or samples. Nat Biotechnol 32, 896-902.

Rosenberg, A.B., Roco, C.M., Muscat, R.A., Kuchina, A., Sample, P., Yao, Z., Graybuck, L.T., Peeler, D.J., Mukherjee, S., Chen, W., et al. (2018). Single-cell profiling of the developing mouse brain and spinal cord with split-pool barcoding. Science.

Rudy, B., Fishell, G., Lee, S., and Hjerling-Leffler, J. (2011). Three groups of interneurons account for nearly 100% of neocortical GABAergic neurons. Dev Neurobiol 71, 45-61.

Saunders, A., Macosko, E.Z., Wysoker, A., Goldman, M., Krienen, F.M., de Rivera, H., Bien, E., Baum, M., Bortolin, L., Wang, S., et al. (2018). Molecular Diversity and Specializations among the Cells of the Adult Mouse Brain. Cell 174, 1015-1030 e1016.

Sorensen, S.A., Bernard, A., Menon, V., Royall, J.J., Glattfelder, K.J., Desta, T., Hirokawa, K., Mortrud, M., Miller, J.A., Zeng, H., et al. (2015). Correlated gene expression and target specificity demonstrate excitatory projection neuron diversity. Cereb Cortex 25, 433-449.

Supek, F., Bosnjak, M., Skunca, N., and Smuc, T. (2011). REVIGO summarizes and visualizes long lists of gene ontology terms. PLoS One 6, e21800.

Tasic, B., Menon, V., Nguyen, T.N., Kim, T.K., Jarsky, T., Yao, Z., Levi, B., Gray, L.T., Sorensen, S.A., Dolbeare, T., et al. (2016). Adult mouse cortical cell taxonomy revealed by single cell transcriptomics. Nature neuroscience 19, 335-346.

Tosches, M.A., Yamawaki, T.M., Naumann, R.K., Jacobi, A.A., Tushev, G., and Laurent, G. (2018). Evolution of pallium, hippocampus, and cortical cell types revealed by single-cell transcriptomics in reptiles. Science 360, 881-888.

Tripodi, R.V., Sebastiano; Pelillo, Marcello (2016). Context Aware Nonnegative Matrix Factorization Clustering. ARXIV.

Tsaytler, P., Harding, H.P., Ron, D., and Bertolotti, A. (2011). Selective inhibition of a regulatory subunit of protein phosphatase 1 restores proteostasis. Science 332, 91-94.

Tsuneoka, Y., Tsukahara, S., Yoshida, S., Takase, K., Oda, S., Kuroda, M., and Funato, H. (2017). Moxd1 Is a Marker for Sexual Dimorphism in the Medial Preoptic Area, Bed Nucleus of the Stria Terminalis and Medial Amygdala. Front Neuroanat 11, 26.

Wang, X., Allen, W.E., Wright, M.A., Sylwestrak, E.L., Samusik, N., Vesuna, S., Evans, K., Liu, C., Ramakrishnan, C., Liu, J., et al. (2018). Three-dimensional intact-tissue sequencing of single-cell transcriptional states. Science 361.

Xiong, Y., Mahmood, A., Meng, Y., Zhang, Y., Zhang, Z.G., Morris, D.C., and Chopp, M. (2012). Neuroprotective and neurorestorative effects of thymosin beta4 treatment following experimental traumatic brain injury. Ann N Y Acad Sci 1270, 51-58.

Xu, W.L., Xin; Gong, Yihong (2003). Document clustering based on non-negative matrix factorization. Paper presented at: ACM SIGIR.

Xu, X., Coats, J.K., Yang, C.F., Wang, A., Ahmed, O.M., Alvarado, M., Izumi, T., and Shah, N.M. (2012). Modular genetic control of sexually dimorphic behaviors. Cell 148, 596-607.

Yang, Z., and Michailidis, G. (2016). A non-negative matrix factorization method for detecting modules in heterogeneous omics multi-modal data. Bioinformatics 32, 1-8.

Zeisel, A., Hochgerner, H., Lonnerberg, P., Johnsson, A., Memic, F., van der Zwan, J., Haring, M., Braun, E., Borm, L.E., La Manno, G., et al. (2018). Molecular Architecture of the Mouse Nervous System. Cell 174, 999-1014 e1022.

Zheng, G.X., Terry, J.M., Belgrader, P., Ryvkin, P., Bent, Z.W., Wilson, R., Ziraldo, S.B., Wheeler, T.D., McDermott, G.P., Zhu, J., et al. (2017). Massively parallel digital transcriptional profiling of single cells. Nat Commun 8, 14049.

